# Human skin specific long noncoding RNA *HOXC13-AS* regulates epidermal differentiation by interfering with Golgi-ER retrograde transport

**DOI:** 10.1101/2022.07.04.498689

**Authors:** Letian Zhang, Minna Piipponen, Zhuang Liu, Dongqing Li, Xiaowei Bian, Maria A. Toma, Pehr Sommar, Ning Xu Landén

## Abstract

After a skin injury, keratinocytes switch from a state of homeostasis to one of regeneration leading to the reconstruction of the epidermal barrier. The regulatory mechanism of gene expression underpinning this key switch during human skin wound healing is enigmatic. Long noncoding RNAs (lncRNAs) constitute a new horizon in the understanding of the regulatory programmes encoded in the mammalian genome. By comparing the transcriptome of an acute human wound and skin from the same donor as well as keratinocytes isolated from these paired tissue samples, we generated a list of lncRNAs showing changed expression in keratinocytes during wound repair. Our study focused on *HOXC13-AS*, a recently evolved human lncRNA specifically expressed in epidermal keratinocytes, and we found that its expression was temporally downregulated during wound healing. In line with its enrichment in suprabasal keratinocytes, *HOXC13-AS* was found to be increasingly expressed during keratinocyte differentiation, but its expression was reduced by EGFR signalling. After *HOXC13-AS* knockdown or overexpression in human primary keratinocytes undergoing differentiation induced by cell suspension or calcium treatment and in organotypic epidermis, we found that keratinocyte differentiation was promoted, while the cell inflammatory response was suppressed. Moreover, RNA pull-down assays followed by mass spectrometry and RNA immunoprecipitation analysis revealed that mechanistically *HOXC13-AS* sequestered the coat complex subunit alpha (COPA) protein and interfered with Golgi-to-endoplasmic reticulum (ER) molecular transport, resulting in ER stress and enhanced keratinocyte differentiation. In summary, we identified *HOXC13-AS* as a crucial regulator of human epidermal differentiation.

## INTRODUCTION

The epidermis is the outermost epithelium layer and is stratified, and it protects the human body from external stimuli and prevents dehydration^1^. Keratinocytes constitute approximately 90% of all epidermal cells and undergo terminal differentiation to form the basal, spinous, granular, and cornified layers of the epidermis. Keratinocyte differentiation has been characterized by a dynamically changed gene expression programme, e.g., early differentiated keratinocytes express KRT1, KRT10, and IVL, and then, LOR and FLG levels increase at later differentiation stages^2, 3^. Well-balanced keratinocyte proliferation and differentiation are essential for maintaining epidermal homeostasis, which is disrupted by skin injury, and wound-edge keratinocytes swiftly switch their status to engage in regeneration^4^. The dynamic gene expression and related regulatory mechanisms underpinning the switch between keratinocytes in the epidermal homeostasis state and the regeneration state are not fully understood; The mechanism is even more elusive in the human tissue environment during wound healing. Addressing fundamental questions about homeostasis-to-regeneration phenotype switching is required to understand the pathological mechanism underlying failed re-epithelization in chronic nonhealing wounds, which have led to major and increasing health and financial burdens worldwide^5, 6^.

In addition to protein-coding genes, most of the human genome comprises a vast landscape of regulatory elements, including tens of thousands of long noncoding RNAs (lncRNAs). LncRNAs are transcripts longer than 200 nucleotides that with no or limited sequences that can be translated^7, 8^. Increasing numbers of lncRNAs have been shown to regulate vital cellular processes via a large variety of molecular mechanisms and to play critical roles in health and disease^9^. Importantly, both the expression pattern of lncRNAs and their functions are more cell-type- and cell-state-specific than protein-coding genes, which endows lncRNAs with promising therapeutic and diagnostic potential^10–12^. In the skin, a few lncRNAs, including *ANCR*, *TINCR*, *LINC00941*, *uc.291,* and *PRANCR*, have been shown to regulate keratinocyte differentiation^13–17^. Moreover, three lncRNAs have been reported to function in keratinocytes during skin wound healing, i.e., *WAKMAR2* suppresses the inflammatory response, while *WAKMAR1*, *WAKMAR2*, and *TETILA* change cell mobility^18–20^. Although in its infancy, this field has produced evidence suggesting that lncRNAs are important regulators in epidermal homeostasis and regeneration, and further efforts to gain a more holistic and deeper understanding of these RNAs are warranted.

In this study, by tracing the *in vivo* transcriptomic changes in keratinocytes during human skin wound healing, we generated a list of lncRNAs that changed during the switch between the epidermal homeostasis state and the regeneration state. We focused on a human skin- and keratinocyte-specific lncRNA, *HOXC13-AS*, and revealed the crucial role it plays in regulating keratinocyte differentiation by interfering with retrograde protein transport from the Golgi to the endoplasmic reticulum (ER). The temporal downregulation of *HOXC13-AS* in wound-edge keratinocytes, likely due to high EGFR signalling during wound repair, and its restored expression during re-epithelialization reflect its physiological importance in the maintenance and reconstruction of the epidermal barrier.

## RESULTS

### Downregulation of *HOXC13-AS* expression in human wound-edge keratinocytes

To characterize the role played by lncRNAs in human skin wound healing, we created wounds on the skin of healthy donors and collected the wound-edge tissues on day one (acute wound day one, AW1), seven (AW7), and 30 (AW30) until the wounds closed (Fig. 1a). With ribosomal RNA-depleted long RNA sequencing (RNA-seq) of these full-thickness tissue biopsy samples, we identified 57 upregulated and 211 downregulated lncRNAs on AW7 compared to the expression of these lncRNAs in matched skin from the five donors [|fold change| >= 2, False discovery rate (FDR) < 0.05, Fig. 1b, Supplementary Data 1]. To detect keratinocyte-related lncRNA expression changes, we conducted RNA-seq with epidermal CD45-cells and identified 15 upregulated and 671 downregulated lncRNAs in keratinocytes isolated from the AW7 wound edges compared to expression in the matched skin samples (|fold change| >= 2, FDR < 0.05, Fig. 1b, Supplementary Data 2). Interestingly, 3 upregulated and 16 downregulated lncRNAs were identified in both the tissue and epidermal cell RNA-seq analyses (Fig. 1b). We ranked these 19 lncRNAs on the basis of their skin specificity scores, which were calculated by the SPECS method^21^ with the RNA-seq data obtained from 31 human normal tissues in the GTEx database^22^. HOXC13 antisense RNA (*HOXC13-AS*) was identified as a skin-specific lncRNA, with a score even higher than some known skin-specific genes, e.g., *FLG* and *KRT10* (Fig. 1c, d). The highly specific expression pattern of *HOXC13-AS* suggests its potentially unique function in skin, which prompted us to take a closer look at this lncRNA.

**Figure 1.**
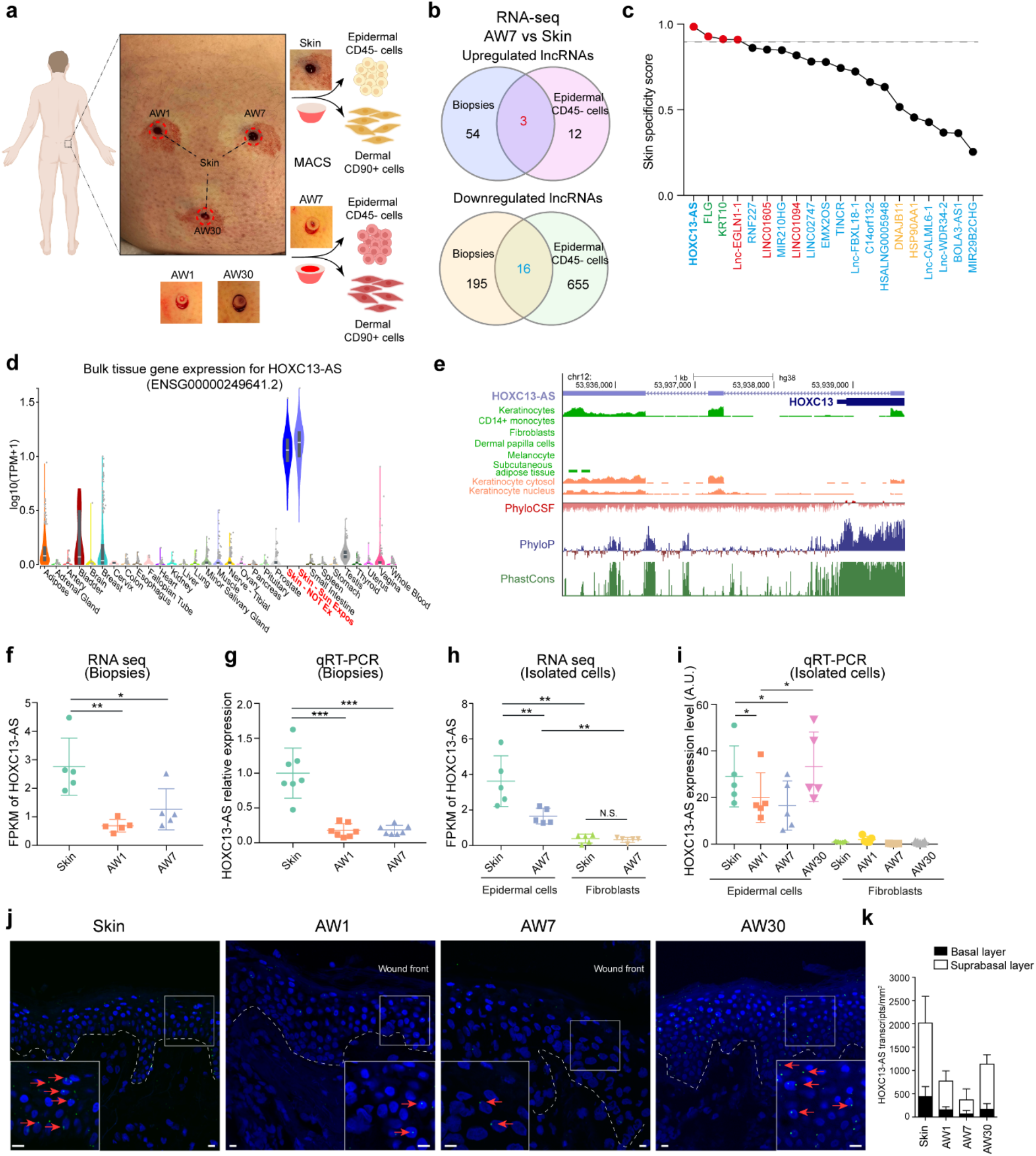
Downregulation of *HOXC13-AS* expression in human wound-edge keratinocytes. **a** Human *in vivo* wound model: full-thickness excisional wounds were created on the skin of healthy volunteers and wound-edge tissues were collected 1 (AW1), 7 (AW7), and 30 days later (AW30) from the same donor. Epidermal CD45^-^ cells (enriched with keratinocytes) and dermal CD90^+^ cells (fibroblasts) were isolated from the skin and AW7 biopsies by Magnetic-Activated Cell Sorting (MACS). RNA-sequencing was performed in both the tissues and the cells. **b** Venn diagram showing the differentially expressed lncRNAs in the CD45^-^ epidermal cells and the tissue biopsies of the AW7 compared to the skin (|Fold change| >= 2, p < 0.05). **c** Skin specificity scores of the differentially expressed lncRNAs surfaced in both the tissue and the epidermal cell RNA-seq analysis (upregulated in red, downregulated in blue), known skin-specific genes KRT10 and FLG (green), and broadly expressed genes DNAJB11 and HSP90AA1 (yellow). The dashed line indicates the skin specificity score of 0.9. **d** *HOXC13-AS* expression data across 31 normal human tissues retrieved from the GTEx database. **e** Genomic snapshot of *HOXC13-AS* generated in GENECODE V38. Data were retrieved from Encyclopedia of DNA Elements data hub, phylogenetic information-based codon substitution frequency (PhyloCSF), and conservation tracks (PhyloP and PhastCons). RNA-seq of *HOXC13-AS* in tissue biopsies (n = 5 donors) **f** and isolated cells (n = 5 donors) **h**. Data are normalized as Fragments per kilobase of a transcript, per million mapped reads (FPKM). QRT-PCR analysis of *HOXC13-AS* in tissue biopsies (n = 7 donors) **g** and isolated cells (n = 5 donors) **i**. Representative photographs **j** and quantification **k** of *HOXC13-AS* fluorescence in situ hybridization (FISH) in human skin and wounds (n = 2 donors). Cell nuclei were co-stained with DAPI (scale bar = 10 μm). \**P* < 0.05; \*\**P* < 0.01; \*\*\**P* < 0.001 by Mann Whitney test **f**-**h** and paired two-tailed Student’s t test **i**. Data are presented as mean ± SD.

*HOXC13-AS* is a divergent lncRNA transcribed from the opposite strand of the protein-coding gene *HOXC13* on human chromosome 12 [GRCh38/hg38, chr12:53,935,328-53,939,643] (Fig. 1e, Supplementary Fig. 1a). It is a recently evolved human lncRNA, as indicated by the low PhyloP and PhastCons scores, and no homologue in rodents has been identified. Moreover, a phylogenetic information-based codon substitution frequency analysis (PhyloCSF) suggested that *HOXC13-AS* lacks protein-coding potential, which was in agreement with a coding potential calculator CPC2 analysis^23^ showing that the coding potential of *HOXC13-AS* was even lower than that of *HOTAIR*^24^, a well-known lncRNA (Supplementary Fig. 1b).

Our RNA-seq analysis of human wounds revealed that *HOXC13-AS* expression was downregulated by AW1 and remained low through AW7 compared to that in the matching skin, and these findings were confirmed by real-time RT–PCR (qRT–PCR) analysis of tissue biopsy samples from seven other donors (Fig. 1f, g). Moreover, we performed RNA-seq and qRT–PCR with paired epidermal keratinocytes and dermal fibroblasts isolated from skin and wound samples on AW7 from 10 healthy donors. We found that in human skin, *HOXC13-AS* was mainly expressed in keratinocytes but not in fibroblasts (Fig. 1h, i), which was in line with the public ENCODE data regarding *HOXC13-AS* expression in different cell types (Fig. 1e). Keratinocyte *HOXC13-AS* expression was transiently downregulated on AW1 and AW7 and then recovered to the level in paired skin on AW30, at which point the wounds had re-epithelized and showed epidermal stratification (Fig. 1i). Performing fluorescence in situ hybridization (FISH), we not only confirmed these findings but also localized *HOXC13-AS* mainly in the suprabasal layers of the epidermis (Fig. 1j, k). Additionally, in published RNA-seq datasets of human wounds, we found reduced *HOXC13-AS* expression in diabetic foot ulcers compared to diabetic foot skin^25^ and on day two and five in acute wounds compared to the expression in the skin^26^ (Supplementary Fig. 1c, d). Consistent with its skin-specific expression (Fig. 1c, d), *HOXC13-AS* was not detected in human oral mucosal wounds^26^ (Supplementary Fig. 1d).

In summary, we identified a human skin keratinocyte-specific lncRNA: *HOXC13-AS*. The expression of this lncRNA was reduced upon skin injury but was restored during re-epithelization, suggesting that *HOXC13-AS* may be involved in the maintenance and reconstruction of the epidermal barrier.

### Single cell transcriptomic analysis of human skin revealed increased *HOXC13-AS* expression in granular keratinocytes

We performed a single-cell RNA sequencing (scRNA-seq) analysis of human skin (n=3) using 10X Chromium technology. Unsupervised clustering of 11800 cells with the Seurat package revealed 21 cell clusters, which included keratinocytes (basal, spinous, and granular keratinocytes), fibroblasts (FB-I − IV), melanocytes (MELs), Schwann cells, pericytes-vascular smooth muscle cells (PC-vSMCs), lymphatic and vascular endothelial cells (LE and VE), and immune cells [mast cells, NK cells, B cells, monocytes-macrophages (Monos-Macs), Langerhans cells (LC), and dendritic cells (DC)] (Fig. 2a). In line with the bulk RNA-seq data obtained with individual cell types (Fig. 1e, h), scRNA-seq demonstrated keratinocyte-specific *HOXC13-AS* expression in human skin (Fig. 2a). Interestingly, we found that *HOXC13-AS* was predominantly expressed in granular keratinocytes (Fig. 2b), which agreed with our FISH results showing higher *HOXC13-AS* expression in the suprabasal layers of the epidermis (Fig. 1j, k). Additionally, a pseudotime analysis showed that all keratinocytes were ordered along a differentiation trajectory, which revealed that *HOXC13-AS* expression increased with keratinocyte differentiation (Fig. 2c). Furthermore, to envisage the potential functional role of *HOXC13-AS*, we performed an expression correlation analysis between *HOXC13-AS* and all other genes expressed in granular keratinocytes, and thus, we identified 1935 positively correlated genes (R > 0, *P* < 0.05). Gene Ontology analysis (GO) unravelled that the 50 genes that were most correlated with *HOXC13-AS* were mainly involved in keratinocyte differentiation and the immune response, suggesting that this lncRNA may play a similar role in these pathways (Fig. 2d, Supplementary Data 3).

**Figure 2.**
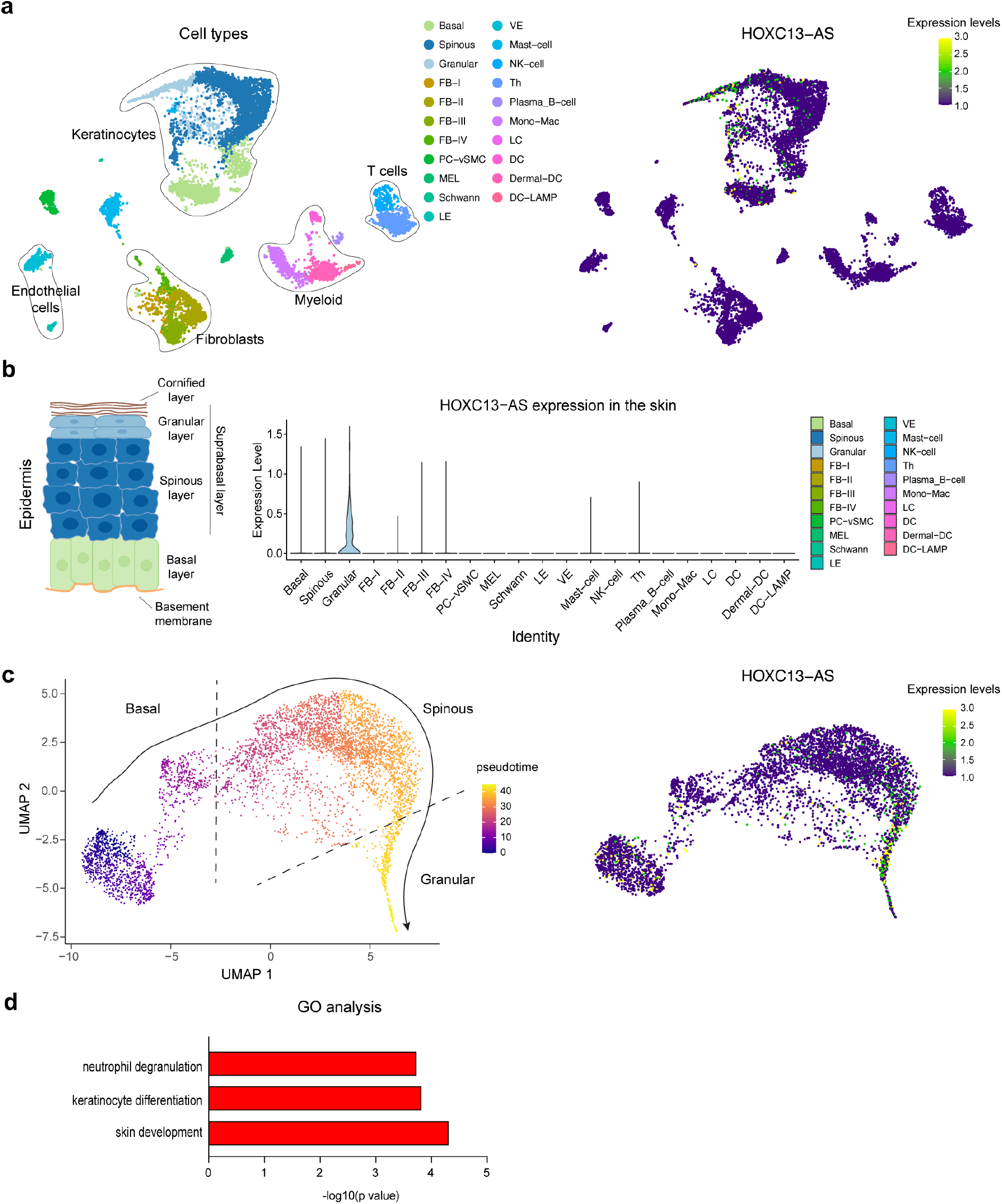
Granular keratinocytes are the major cells expressing *HOXC13-AS* in the skin. **a** UMAP representation of all cell types (left) and *HOXC13-AS* expressing cells (right) identified in human skin (n = 3 donors) by single cell RNA-sequencing (scRNA-seq). **b** Violin plots of *HOXC13-AS* expression in different cell types in human skin. **c** Pseudotime trajectory of all the keratinocytes (i.e., basal, spinous, and granular keratinocytes) colored by pseudotime (left) and *HOXC13-AS* expression (right). **d** Gene Ontology (GO) analysis of the top 50 genes with expression positively correlated with *HOXC13-AS* in granular keratinocytes analyzed by scRNA-seq.

### *HOXC13-AS* expression is regulated in the opposite direction by keratinocyte growth and differentiation signalling

We next explored the mechanism that modulates *HOXC13-AS* expression in epidermal homeostasis and regeneration. We treated human keratinocytes with a panel of growth factors (VEGF-A, FGF2, EGF, KGF, HB-EGF, IGF-1, TGF-β1, and TGF-β3) and cytokines (IL-1α, IL-6, IL-23, IL-36α, TNFα, MCP-1, and GM-CSF) known to be important to wound repair^27^. A qRT-PCR analysis revealed that *HOXC13-AS* expression was significantly downregulated by two members of the EGF family (HB-EGF and EGF)^27^ and KGF (Fig. 3a, Supplementary Fig. 2a). Additionally, reduced *HOXC13-AS* expression was found in a public dataset (GSE156089) generated with an RNA-seq analysis of epidermal stem cells treated with EGF (Supplementary Fig. 2b). Moreover, we showed that blocking EGF receptor (EGFR) signalling with the chemical inhibitor AG1478^28^ prominently enhanced *HOXC13-AS* expression in keratinocytes in a dose-dependent manner, which confirmed the inhibitory effect of growth factor signalling on *HOXC13-AS* expression in keratinocytes (Fig. 3b). Importantly, we found that the expression of HB-EGF and KGF was significantly upregulated during human skin wound repair and a significantly negative correlation between *HOXC13-AS* and *HBEGF* expression, as shown by the RNA-seq analysis of human wound tissues and isolated cells, suggesting that enhanced growth factor signalling may contribute to the downregulation of *HOXC13-AS* expression in human wounds *in vivo* (Fig. 3c, d, Supplementary Fig. 2c).

**Figure 3.**
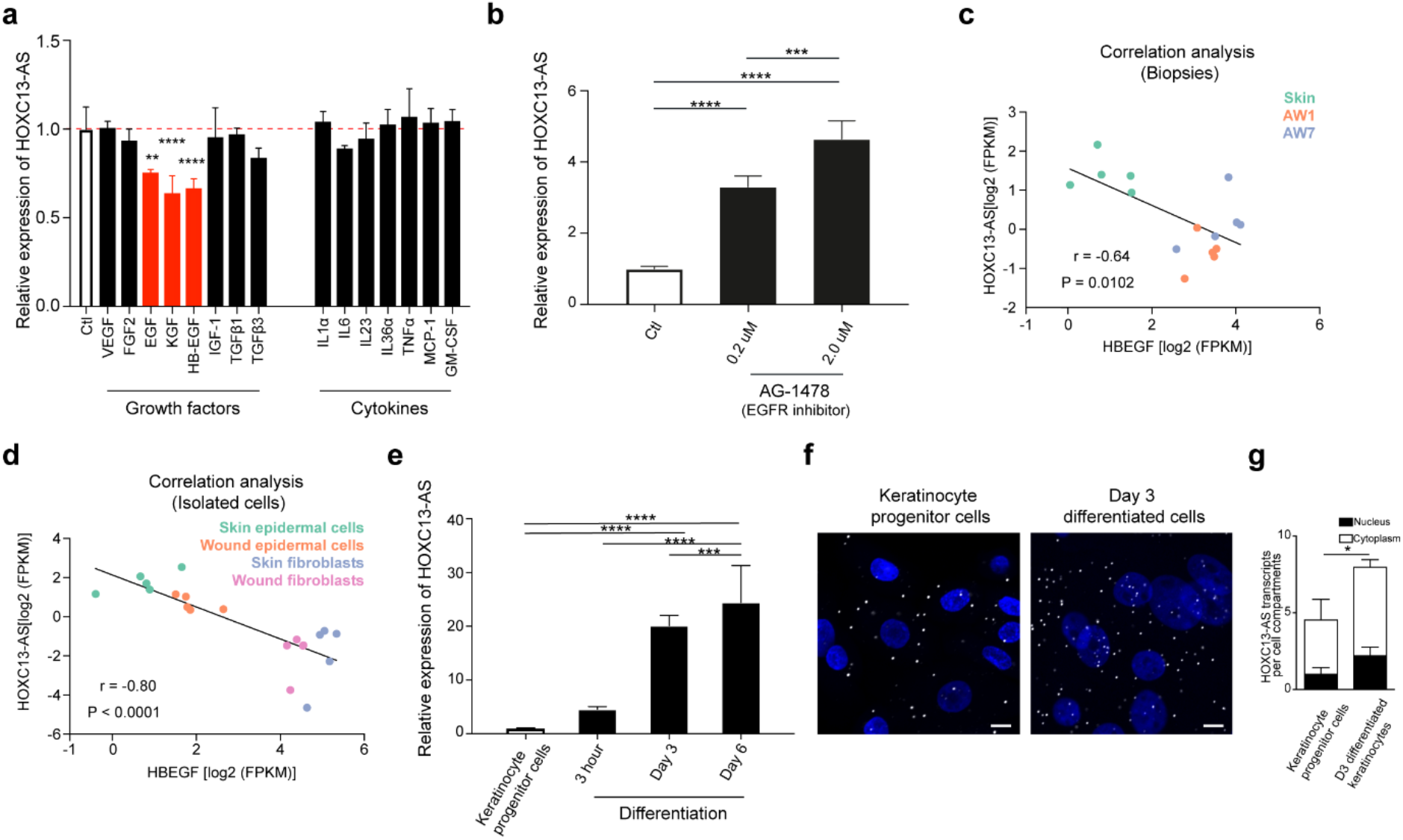
*HOXC13-AS* expression is regulated in the opposite direction by keratinocyte growth and differentiation signalling. QRT-PCR analysis of *HOXC13-AS* expression in keratinocytes treated with wound-related cytokines, growth factors **a,** or AG-1478 **b** for 24 hours (n = 4). Expression correlation of *HOXC13-AS* with *HBEGF* in the skin and wound tissues **c** and the isolated cells **d** analyzed by RNA-seq. **e** qRT-PCR analysis of *HOXC13-AS* in keratinocyte progenitor cells and calcium-induced differentiated keratinocytes (n = 4). Representative photographs **f** and quantification **g** of *HOXC13-AS* FISH in keratinocyte progenitor cells and differentiated keratinocytes treated with 1.5 mM calcium for 3 days (n = 3). Cell nuclei were co-stained with DAPI. Scale bar = 20 μm. \**P* < 0.05; \*\**P* < 0.01, \*\*\**P* < 0.001 and \*\*\*\**P* < 0.0001 by unpaired two-tailed Student’s t test **a**, **b**, **e**, **g**, or Pearson’s correlation test **c** and **d**. Data are presented as mean ± SD.

As the FISH and scRNA-seq analyses of human skin showed higher *HOXC13-AS* expression in more-differentiated keratinocytes (Fig. 1j, k and Fig. 2c) and because EGFR signalling inhibition has been reported to induce keratinocyte differentiation^26^, we next examined whether keratinocytes undergoing differentiation showed enhanced *HOXC13-AS* expression. Studying a calcium-induced keratinocyte differentiation model^29, 30^, we found that *HOXC13-AS* expression gradually increased as cell differentiation progressed, as shown by both *HOXC13-AS* qRT–PCR and FISH analysis results (Fig. 3e-g, Supplementary Fig. 2d).

Considering these data, we concluded that *HOXC13-AS* expression in keratinocytes was induced by cell differentiation but suppressed by growth factor signalling, which explains its increased expression in the suprabasal layers of the epidermis that contain more differentiated keratinocytes and its decreased expression during wound repair due to high growth signalling (Fig. 1 and Fig. 2).

### *HOXC13-AS* promotes keratinocyte differentiation

To characterize the potential functional role played by *HOXC13-AS*, we knocked down (KD) *HOXC13-AS* expression by transfecting human keratinocytes with *HOXC13-AS*-specific short interfering RNAs (siRNAs), and the loss of *HOXC13-AS* was confirmed by FISH (Fig. 4a, c) and qRT–PCR analyses (Supplementary Fig. 3a). In addition, we overexpressed (OE) *HOXC13-AS* in keratinocytes using a *HOXC13-AS* expression vector (pcDNA-*HOXC13-AS*), which significantly increased *HOXC13-AS* levels in the keratinocytes (Fig. 4b, c, Supplementary Fig. 3b). We showed that neither *HOXC13-AS* KD nor OE affected *HOXC13* expression in progenitor or differentiated keratinocytes (Supplementary Fig. 3c-e).

**Figure 4.**
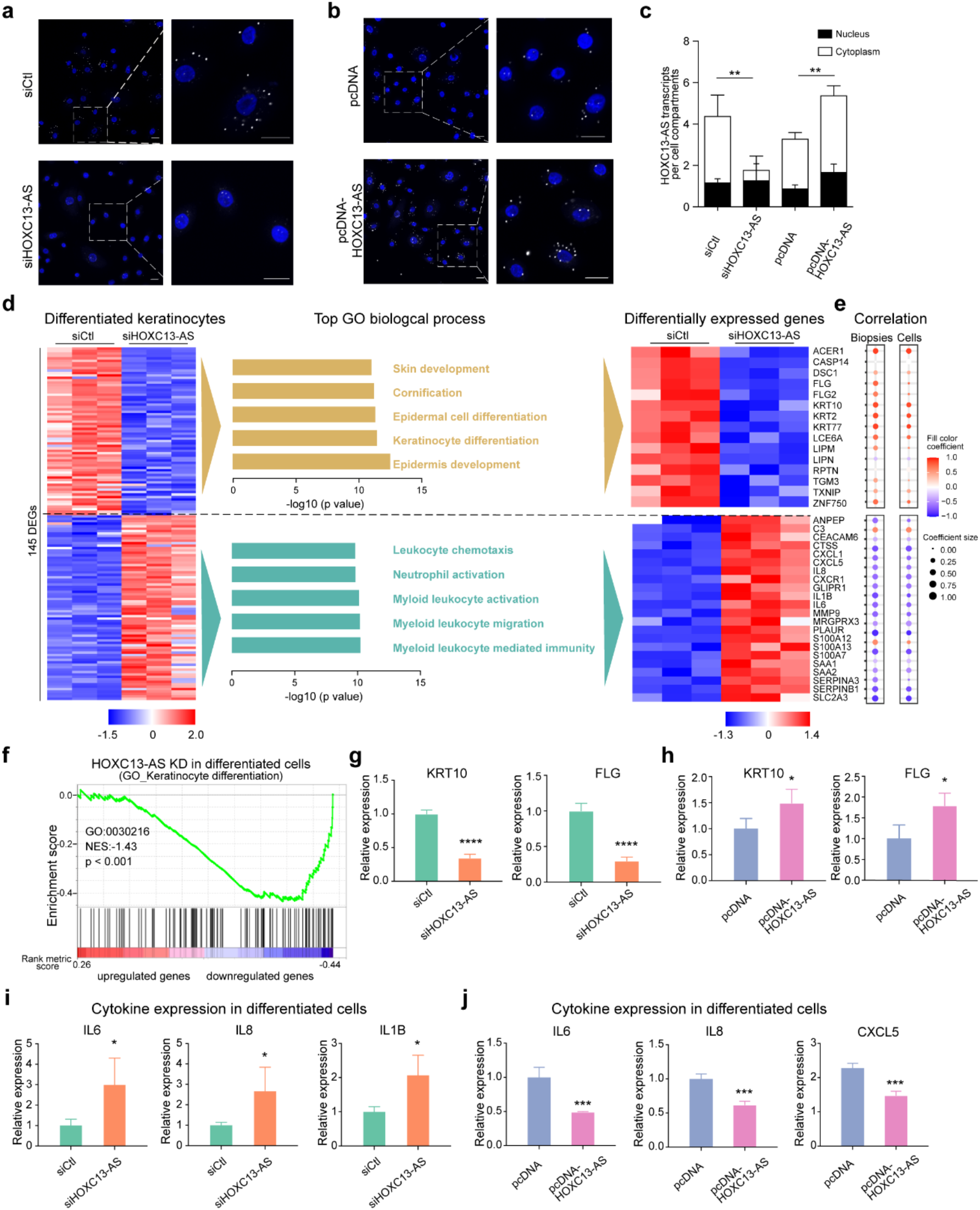
*HOXC13-AS* regulates keratinocyte differentiation and inflammatory response. Representative photographs **a, b** and quantification **c** of *HOXC13-AS* FISH in keratinocytes transfected with *HOXC13-AS* siRNA pool or nontargeting control (siCtl) **a**, pcDNA-*HOXC13-AS* or empty vector **b** (n= 3). Cell nuclei were co-stained with DAPI. Scale bar = 20 μm. **d** Microarray analysis of differentiated keratinocytes transfected with *HOXC13-AS* siRNA pool or siCtl. Heatmap (left panel) illustrates the differentially expressed genes (DEGs, |Fold change| >= 2, FDR < 0.05). GO analysis of the DEGs is shown in the middle panel. The DEGs associated with the GO terms are shown in the heatmap (right panel). **e** Their expression correlation with *HOXC13-AS* in the skin and wound biopsies and isolated epidermal cells analyzed by RNA-seq. **f** GSEA evaluated enrichment for the keratinocyte differentiation-related genes (GO:0030216) in the microarray data. NES, normalized enrichment score. qRT-PCR analysis of *KRT10* and *FLG* expression in differentiated keratinocytes transfected with *HOXC13-AS* siRNA pool **g** or pcDNA-*HOXC13-AS* **h** compared to respective controls (n = 4). qRT-PCR analysis of cytokine expression in differentiated keratinocytes transfected with *HOXC13-AS* siRNA pool **i** or pcDNA-*HOXC13-AS* **j** compared to respective controls (n = 4). \**P* < 0.05; \*\**P* < 0.01, \*\*\**P* < 0.001, and \*\*\*\**P* < 0.0001 by unpaired two-tailed Student’s t test **c**, **g** - **j**, or Pearson’s correlation test **e**. Data are presented as mean ± SD.

Next, we performed a microarray analysis with calcium-induced differentiated keratinocytes in which *HOXC13-AS* expression was knocked down, which led to the identification of 76 upregulated and 69 downregulated genes (|fold change| >= 2, p value < 0.05) (Fig. 4d). A GO analysis revealed that the expression of genes involved in keratinocyte differentiation (e.g., *KRT10*, *FLG*, *DSC1*^31^, and *CASP14*^32^) was decreased, whereas the expression of immune response-related genes (e.g., *IL6, IL8,* and *IL1B*) was increased after *HOXC13-AS* was knocked down (Fig. 4d). Furthermore, we evaluated the expression of these *HOXC13-AS*-regulated genes in our RNA-seq datasets of human wound tissues and isolated epidermal cells (Fig. 1a). We found that differentiation- and inflammation-related genes were positively and negatively correlated with *HOXC13-AS* levels, respectively, supporting the *in vivo* relevance of our findings to *HOXC13-AS-*mediated gene regulation (Fig. 4e, Supplementary Fig. 3f-i). Additionally, a gene set enrichment analysis (GSEA)^33^ confirmed that among the downregulated genes after *HOXC13-AS* was knocked down, keratinocyte differentiation-related genes (GO: 0030216) were significantly (p < 0.001) enriched (Fig. 4f). Furthermore, we confirmed the microarray findings by performing a qRT–PCR analysis of the keratinocyte differentiation markers *KRT10* and *FLG* and the inflammatory genes *IL6, IL8, IL1B, CXCL5*, *CXCL1*, and *MMP9* in a calcium-induced keratinocyte differentiation model with *HOXC13-AS* overexpressed and knocked down (Fig. 4g-j, Supplementary Fig. 3j). Both gain- and loss-of-function studies suggested that *HOXC13-AS* promoted keratinocyte differentiation while suppressing the cellular inflammatory response. Additionally, we showed that neither knocking down nor overexpressing *HOXC13-AS* affected human progenitor keratinocyte proliferation or migration (Supplementary Fig. 4a-c).

Moreover, we utilized a suspension-induced keratinocyte differentiation model (Fig. 5a)^34, 35^. Human progenitor keratinocytes were cultured in a single-cell suspension from 2 to 24 hours, which induced cell differentiation, as shown by the gradually increased expression of the differentiation markers *KRT10*, *IVL*, and *FLG* (Fig. 5b-d). Additionally, *HOXC13-AS* expression was enhanced through cell differentiation (Fig. 5e). After confirming the high efficiency of *HOXC13-AS* knockdown (KD) and overexpression (OE) in this model (Supplementary Fig. 4d), we showed that *HOXC13-AS* OE led to enhanced keratinocyte differentiation, whereas *HOXC13-AS* KD led to profoundly reduced suspension-induced keratinocyte differentiation, as evidenced by the changes in *KRT10*, *FLG*, and *IVL* at the mRNA (Fig. 5f-h) and protein levels (Fig. 5i).

**Figure 5.**
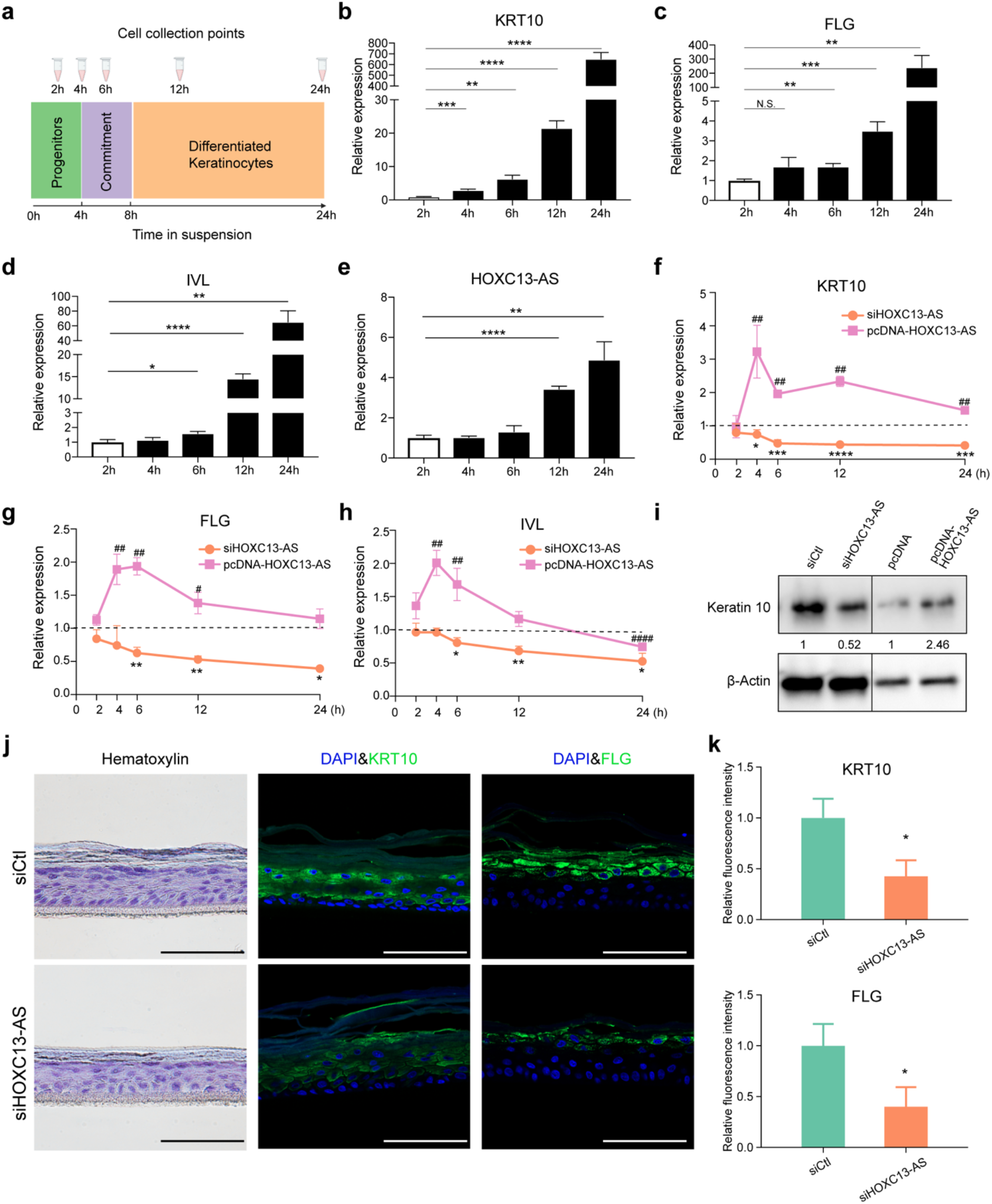
*HOXC13-AS* regulates keratinocyte differentiation in suspension-induced differentiation model and organotypic human epidermal tissues. **a** Schematic representation of the experimental flow (left panel) and **b-e** qRT-PCR analysis of *KRT10*, *FLG*, *IVL,* and *HOXC13-AS* expression in the suspension-induced keratinocyte differentiation model (n = 3). qRT-PCR analysis of *KRT10* **f,** *FLG* **g,** and *IVL* **h** in keratinocytes transfected with *HOXC13-AS* siRNA pool, pcDNA-*HOXC13-AS* followed by suspension-induced differentiation. The results were normalized with the respective controls (n = 3). **i** Western blot of Keratin 10 in keratinocytes transfected with *HOXC13-AS* siRNA pool, pcDNA-*HOXC13-AS,* or respective controls after suspension induction for 24 hours. **j** Representative photograph of hematoxylin and immunofluorescent staining of the organotypic epidermis with *HOXC13-AS* knockdown. Cell nuclei were co-stained with DAPI. Scale bar = 100 μm. **k** Quantification of fluorescence intensities of KRT10 and FLG in organotypic epidermis with *HOXC13-AS* knockdown (n = 3). * or ^#^ *P* < 0.05; ** or ^##^ *P* < 0.01, *** or ^###^ *P* < 0.001 and **** or ^####^ *P* < 0.0001 by unpaired two-tailed Student’s t test **b - h, k**. Data are presented as mean ± SD.

To functionally characterize *HOXC13-AS* in a tissue environment, we generated organotypic human epidermal tissues^16^ using progenitor keratinocytes with or without *HOXC13-AS* knocked down (Supplementary Fig. 4e). By performing immunofluorescence staining (IF), we showed that the abundance of the differentiation markers KRT10 and FLG in suprabasal layer keratinocytes was significantly reduced after *HOXC13-AS* KD, suggesting compromised keratinocyte differentiation in the *HOXC13-AS*-deficient organotypic epidermis (Fig. 5j, k). Notably, in these organotypic tissues, we did not detect apoptotic cells through IF of caspase 3 (Supplementary Fig. 4e). The number of proliferating keratinocytes, which were localized to the basal layer of the epidermis, as indicated by Ki67 IF, was not changed by *HOXC13-AS* KD (Supplementary Fig. 4f).

Collectively, these data obtained with multiple physiologically relevant keratinocyte differentiation models demonstrated a prominent role for *HOXC13-AS* in promoting keratinocyte differentiation and showed that *HOXC13-AS* upregulation is required for maintaining the epidermal barrier.

### *HOXC13-AS* interacts with COPA protein

We next investigated the molecular mechanisms mediating the pro-differentiation function of *HOXC13-AS* in keratinocytes. In this analysis, subcellular localization is usually an indicator of the modes of action of lncRNAs. By performing cell fractionation assays, we showed a greater abundance of *HOXC13-AS* in the cytoplasm than in the nucleus of keratinocytes (Fig. 6a), which was consistent with our FISH analysis of human skin, wound tissues, and keratinocytes (Fig. 1j, Fig. 3f, Fig. 4a, b), as well as the public ENCODE data (Fig. 1e). The cytoplasmic localization of *HOXC13-AS* indicated that it might act via an *in trans* mode^9, 36^.

**Figure 6.**
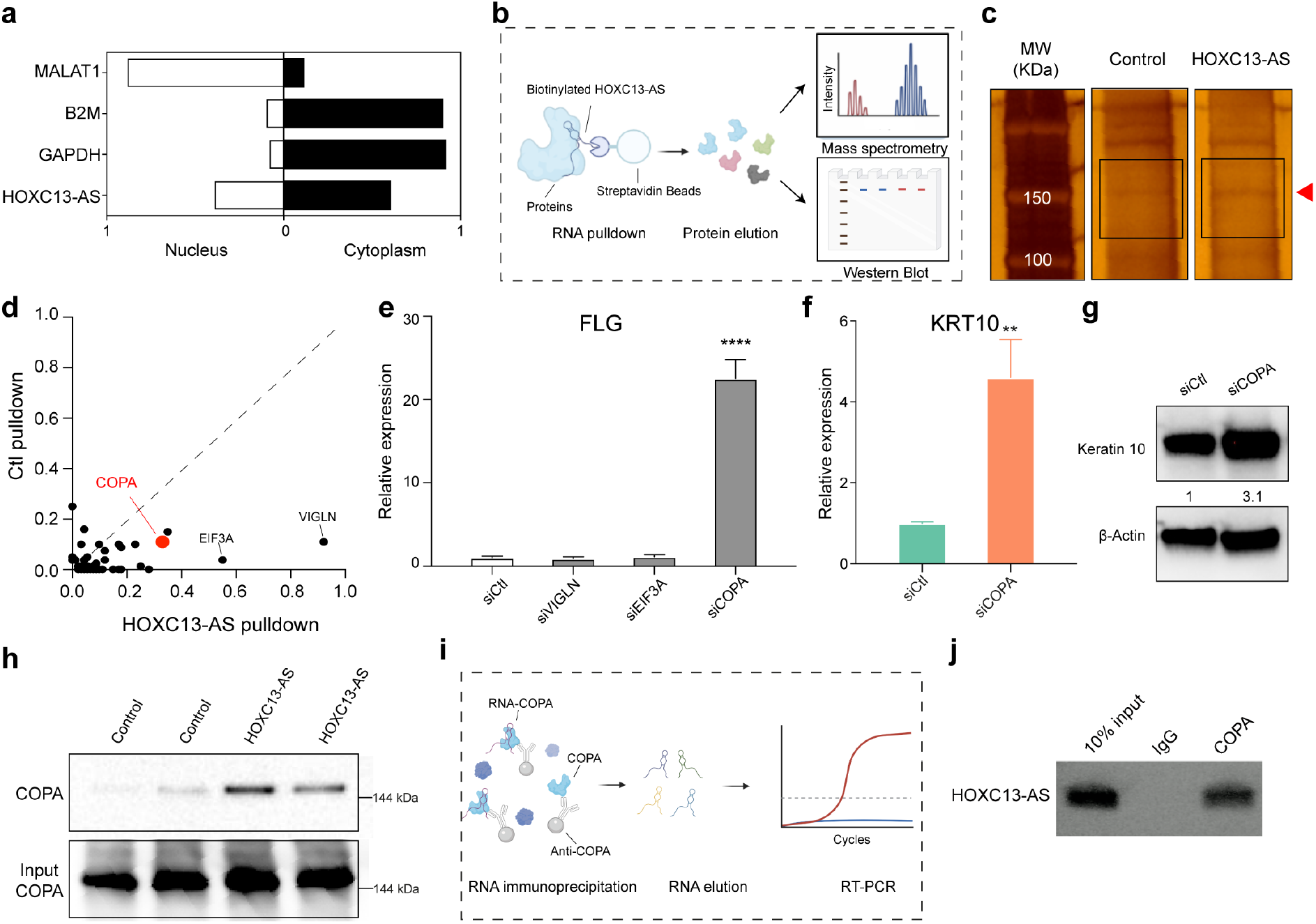
*HOXC13-AS* interacts with COPA protein to regulate keratinocyte differentiation. **a** qRT-PCR analysis of *HOXC13-AS*, *GAPDH*, *B2M,* and *MALAT1* in the nuclear or cytoplasmic fractions of keratinocytes. **b** Schematic representation of the RNA pulldown experiment. **c** Silver staining of proteins bound to *HOXC13-AS* or the control Poly(A)_25_ RNA. The red arrow indicates the differential band, and the rectangles specifies the gel regions sent for mass spectrometry analysis (MS). **d** *HOXC13-AS* bound proteins identified by MS. **e** qRT-PCR analysis of *FLG* expression in differentiated keratinocytes with VIGLN, EIF3A, or COPA expression silencing (n = 3). qRT-PCR **f** and western blot **g** of Keratin 10 expression in differentiated keratinocytes with COPA knockdown (n = 3). **h** Western blot of COPA bound to *HOXC13-AS* or control RNA. Same amount of protein lysates were used as the input. **i** Schematic overview of the RNA immunoprecipitation assays (RIP). **j** Agarose gel electrophoresis of *HOXC13-AS* RT-PCR products with the RNA retrieved from RIP. \*\**P* < 0.01 and \*\*\*\**P* < 0.0001 by unpaired two-tailed Student’s t test **e**, **f**. Data are presented as mean ± SD.

We surveyed the protein interactome of *HOXC13-AS* by performing an RNA pull-down experiment. After incubating keratinocyte protein lysates, biotinylated *HOXC13-AS* together with its binding proteins were purified in process based on streptavidin beads^37^ (Fig. 6b). Running the purified proteins in a gel followed by silver staining, we observed that a band representing approximately 150 kDa was more intense in the *HOXC13-AS* pull-down fraction than in the control poly(A)_25_ RNA pull-down fraction (Fig. 6c). We excised the regions of this differential band and analysed the extract protein by mass spectrometry (MS). High-density lipoprotein binding protein (VIGLN), eukaryotic translation initiation factor 3 subunit A (EIF3A), and coatomer subunit alpha (COPA) were identified as the proteins most enriched in the *HOXC13-AS* pull-down product fraction (Fig. 6d, Supplementary data 4). As *HOXC13-AS* promotes keratinocyte differentiation, we next examined whether any of the identified proteins were involved in the same biological processes. By silencing their expression in keratinocytes with gene-specific siRNAs, we found that only knocking down *COPA* expression significantly changed *FLG* expression in differentiated keratinocytes (Fig. 6e). Furthermore, our qRT–PCR and western blotting analyses revealed that COPA silencing induced *KRT10* expression at both the mRNA and protein levels (Fig. 6f, g). Therefore, our study focused on the COPA protein. We first confirmed the specific pull-down of the COPA protein by *HOXC13-AS* by western blotting with a COPA-specific antibody (Fig. 6h). Next, we performed RNA immunoprecipitation (RIP), which showed that *HOXC13-AS* was pulled down by an anti-COPA antibody but not by IgG (Fig. 6i, j).

In summary, our data suggested that COPA negatively regulates keratinocyte differentiation and that its binding with *HOXC13-AS* may serve as a key link in the mechanisms controlling epidermal differentiation.

### *HOXC13-AS* interferes with Golgi-ER retrograde transport causing ER stress

The COPA protein makes up part of coatomer protein complex I (COPI), which is required for the retrograde transport of cargo proteins from the Golgi to the endoplasmic reticulum (ER) and the movement of vesicles within the Golgi^38^. As *HOXC13-AS* binds *to* COPA protein, we next evaluated whether *HOXC13-AS* affected the COPI-mediated retrograde transport in keratinocytes. To this end, we utilized brefeldin A (BFA), a fungal metabolite that blocks ER-to-Golgi transport^39^, and assessed the extent of the redistribution of the Golgi marker GM130 from the Golgi to the ER^40^. Consistent with previous studies^41^, GM130 IF showed that BFA treatment quickly induced Golgi-ER retrograde transport in keratinocytes (Fig. 7a, b). Interestingly, we found that *HOXC13-AS* KD enhanced and *HOXC13-AS* OE reduced GM130 retrograde transport (Fig. 7c, d and Supplementary Fig. 5a-c).

**Figure 7.**
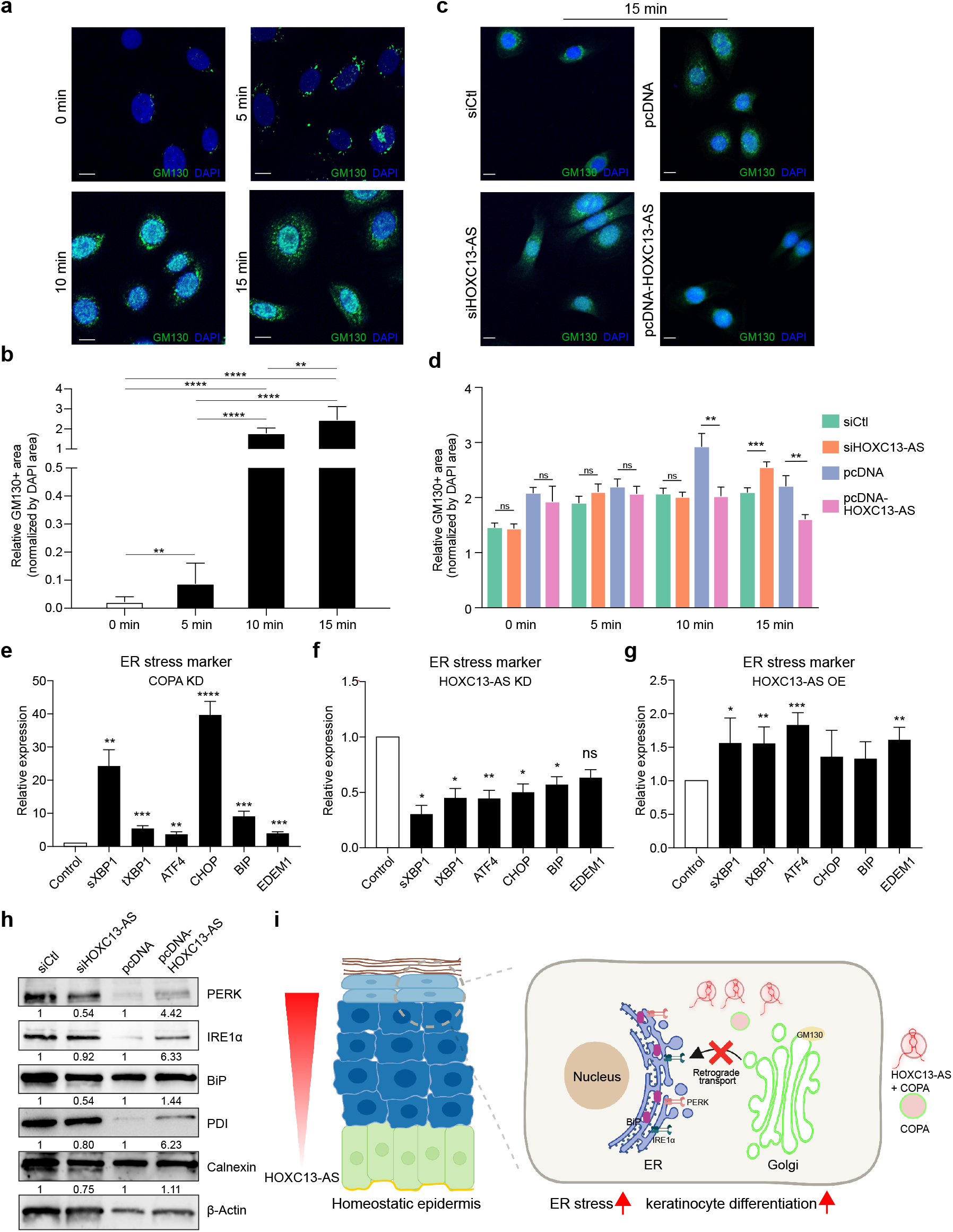
*HOXC13-AS* interferes with Golgi-ER retrograde transport causing ER stress. Representative photograph **a** and quantification **b** of immunofluorescent staining of GM130 in keratinocytes treated with Brefeldin A for 0-15 minutes. Cell nuclei were co-stained with DAPI. (Scale bar = 10 μm, n = 14 - 15 cells). Representative photograph **c** and quantification **d** of immunofluorescent staining of GM130 in keratinocytes transfected with *HOXC13-AS* siRNA pool, pcDNA-*HOXC13-AS* or respective controls, and treated with Brefeldin A. Cell nuclei were co-stained with DAPI (scale bar = 10 μm, n = 8-13 cells at 0 minute and n = 18-22 cells at 5, 10, and 15 minutes). qRT-PCR analysis of ER stress markers in differentiated keratinocytes with COPA knockdown (KD, n = 3) **e**, *HOXC13-AS* KD (n = 4) **f,** or *HOXC13-AS* overexpression (OE, n = 4) **g**. **h** Western blot of ER stress markers in keratinocytes transfected with *HOXC13-AS* siRNA pool, pcDNA-*HOXC13-AS* or respective controls after suspension induction for 24 hours. **i** A proposed model of *HOXC13-AS*/COPA-mediated regulation of keratinocyte differentiation. ns: not significant, \**P* < 0.05, \*\**P* < 0.01, \*\*\**P* < 0.001, and \*\*\*\**P* < 0.0001 by unpaired two-tailed Student’s t test **b**, **d, e** - **g**. Data are presented as mean ± SD **b**, **e** - **g** or mean ± SEM **d**.

COPA mutations have been shown to cause ER stress and to trigger the unfolded protein response (UPR)^42^, which is an adaptive process for maintaining cell viability^43^. Consistent with these previous findings^42^, we showed that COPA silencing induced the expression of ER stress genes, such as activating transcription factor 4 (*ATF4*), total XBP1 (*tXBP1*), spliced XBP1 (*sXBP1*)^44–46^, C/EBP-homologous protein (*CHOP*), binding immunoglobulin protein (*BIP*)^47^, and ER degradation enhancing alpha-mannosidase-like protein 1 (*EDEM1*)^48^, in differentiated keratinocytes (Fig. 7e). Moreover, we found that *HOXC13-AS* KD reduced ER stress gene expression in calcium-induced differentiated keratinocytes, whereas *HOXC13-AS* OE increased ER stress gene expression in these cells (Fig. 7f, g). Similarly, a western blot analysis of a panel of well-established ER stress indicators, including protein kinase RNA-like ER kinase (PERK), inositol-requiring protein 1α (IRE1α), and the chaperone proteins BIP, protein disulfide isomerase (PDI), and calnexin^43, 49, 50^, revealed that *HOXC13-AS* KD inhibited and *HOXC13-AS* OE promoted the accumulation of these ER stress proteins in suspension-induced differentiated keratinocytes (Fig. 7h). Previous studies have shown that physiological ER stress is required for the modulation of keratinocyte differentiation^51^. Here, we demonstrated COPA to be a negative regulator of cell differentiation (Fig. 6e-g). This constellation of findings suggests that by trapping COPA proteins, *HOXC13-AS* hampers Golgi–ER retrograde transport and leads to ER stress, thus promoting keratinocyte differentiation (Fig. 7i).

## DISCUSSION

The search for gene expression regulatory mechanisms driving keratinocyte state switching between homeostasis and regeneration led to our identification of *HOXC13-AS*, a recently evolved, nonconserved human lncRNA. The epidermal-specific expression of *HOXC13-AS* raised the possibility that it may have evolved to regulate epidermis-related functions. As shown in our study, *HOXC13-AS* plays a crucial role in promoting keratinocyte differentiation, leading to the establishment of an effective epidermal barrier. To the best of our knowledge, this is the first report showing the physiological role played by *HOXC13-AS*. This lncRNA has been previously identified as a cancer biomarker in cervical cancer^52^, breast cancer^53^, glioma^54^, hepatocellular carcinoma^55^, cholangiocarcinoma^56^, and head and neck squamous cell carcinoma^57–59^, which emphasizes the absence of *HOXC13-AS* expression in most normal tissues except the skin. Additionally, *HOXC13-AS* reportedly functions as an oncogene promoting proliferation and the epithelial-mesenchymal transition of cancer cells^57^. However, we did not observe a significant effect on normal keratinocyte migration or proliferation (Supplementary Fig. 4a-c). Importantly, our study revealed the highly specific expression and functional pattern of *HOXC13-AS* in human skin, which is an important feature endowing lncRNAs with promising therapeutic and diagnostic potential^10–12^.

Our study adds *HOXC13-AS* to the short list of lncRNAs known to modulate epidermal cell differentiation^13–16^, which includes both negative (*ANCR* and *LINC00941) and positive regulators (TINCR*, *uc.291,* and *PRANCR)*^13–17^. Among these lncRNAs, only *TINCR,* similar to *HOXC13-AS,* was found to be downregulated in human wound tissues and wound-edge keratinocytes compared to the skin in our RNA-seq analysis, (Fig. 1b, c). Prior studies have shown that *TINCR* directly binds to the STAU1 protein, thus stabilizing differentiation-related mRNAs^14^. In addition to acting as a lncRNA, *TINCR* encodes a protein named TINCR-encoded ubiquitin-like protein (TUBL), which promotes keratinocyte proliferation and wound repair^60, 61^. Further study to determine whether TINCR may play a role in keratinocyte homeostasis-to-regeneration state transition is warranted, and if TINCR is found to be involved, the determination of whether it functions as a lncRNA or TUBL in this process will be needed.

We discovered that *HOXC13-AS* acts via a unique mechanism, i.e., sequestration of COPA, a protein required for retrograde Golgi-to-ER transport to recycle the ER-derived transport machinery and resident proteins^62, 63^. Mutant COPA has been previously shown to impair the assembly of proteins targeted for transport and to lead to ER stress and UPR activation in hereditary autoimmune-mediated lung disease and arthritis^38, 42^. Interestingly, several lines of evidence have linked ER stress-triggered UPR with keratinocyte differentiation^64, 65^; e.g., the expression of ER stress and UPR activation markers (sXBP1, CHOP, and BiP) was increased during keratinocyte differentiation, and specific pharmacological ER stressors induced differentiation-related gene expression in a XBP1-dependent mechanism^51^. Our experiment in which we silenced COPA expression in human keratinocytes not only established the importance of COPA in maintaining ER homeostasis (Fig. 7e) but also demonstrated its inhibitory role on keratinocyte differentiation (Fig. 6i, j).

Moreover, our study revealed a novel regulatory mechanism for COPA-mediated Golgi-to-ER retrograde transport. We showed that the lncRNA *HOXC13-AS* traps COPA protein, thus hindering COPA-mediated cargo assembly, which results in ER stress and promotes keratinocyte differentiation (Fig. 7i). In contrast to the physiological levels of ER stress that are required for modulation of keratinocyte differentiation, persistent or excessive levels of ER stress lead to cell death and apoptosis, which has been detected in skin diseases with aberrant epidermal differentiation, such as Darier’s disease^65^. It is tempting to explore whether *HOXC13-AS* may play a pathological role or even serve as a therapeutic target in human diseases with chronic ER stress.

During human skin wound healing, the expression of *HOXC13-AS* in wound-edge keratinocytes was transiently downregulated, which was likely due to high EGFR signalling in the wound environment. It has been previously shown that sustained activation of EGFR signalling suppressed keratinocyte differentiation, whereas its blockade induced differentiation through the activation of Notch signalling^28^. In this study, we added an additional mechanistic link between EGFR signalling and epidermal differentiation, i.e., EGFR signalling inhibited the expression of *HOXC13-AS,* a crucial positive regulator of keratinocyte differentiation. In addition to its differentiation-promoting function, *HOXC13-AS* suppressed the innate immune response in keratinocytes; thus, its quick downregulation upon skin injury may facilitate the initiation of the inflammatory stage of wound repair (Supplementary Fig. 3k). Moreover, in homeostatic skin, the enrichment of *HOXC13-AS* in differentiated keratinocytes, which comprise the outermost layers of the epidermis, may contribute to the maintenance of immune tolerance of the skin barrier, which is constantly exposed to the harsh external environment.

Taken together, the data obtained through our study shows that a human-specific lncRNA, *HOXC13-AS,* is a crucial regulator of epidermal differentiation and that it functions by sequestrating the COPA protein and interfering with Golgi-to-ER transport. These findings contribute to the understanding of the molecular mechanisms required to maintain and regenerate the epidermal barrier. The specific expression in human skin and the critical function in regulating ER stress make HOXC13-AS a potential therapeutic target for a range of cutaneous diseases characterized by chronic ER stress.

## METHODS

### Human *in vivo* wound healing model

Healthy donors (Supplementary Table 1) were recruited and the surgical procedures were conducted at Karolinska University Hospital (Stockholm, Sweden). Two or three full-thickness excisional wounds were made on the skin of each donor by using a 4 mm biopsy punch, and the excised skin was collected as baseline control (Skin). The wound-edge tissue was collected by using a 6 mm biopsy punch one (AW1), seven (AW7), and thirty days later (AW30) (Fig. 1a). Local lidocaine injection was used for anesthesia while sampling. Written informed consent was obtained from all the donors for collecting and using clinical samples. The study was approved by the Stockholm Regional Ethics Committee (Stockholm, Sweden) and conducted according to the Declaration of Helsinki’s principles.

### Cell culture and treatments

Human adult epidermal keratinocytes were cultured in EpiLife medium supplemented with 10% Human Keratinocyte Growth Supplement (HKGS) and 1% Penicillin and Streptomycin at 37℃ in 5% CO_2_ (all from ThermoFisher Scientific, Waltham, MA). Calcium concentration was increased to 1.5 mM in the medium to induce cell differentiation. To study the mechanisms regulating *HOXC13-AS* expression, we treated keratinocytes with EGFR inhibitor AG-1478 (Millipore, Burlington, MA), cytokines and growth factors (Supplementary Table 2), or PBS as a control for 24 hours, and then *HOXC13-AS* expression was analyzed by qRT-PCR. To study the biological function of *HOXC13-AS*, we transfected the third passage of keratinocytes at 60-70% confluence with a 10 nM siRNA pool targeting *HOXC13-AS* or negative control siRNAs (Supplementary Table 3) for 24 hours with Lipofectamine™ 3000 (ThermoFisher Scientific). To identify the function of *HOXC13-AS*-bound proteins, we transfected keratinocytes with 20 nM siRNAs targeting COPA, HDLBP, EIF3A, or negative control siRNAs for 24 hours (Supplementary Table 3).

### Migration and proliferation assays

Human primary keratinocytes were plated in Essen ImageLock 96-well plates (Sartorius, Göttingen, Germany) at 15,000 cells per well for migration assay and in 12-well plates at 20,000 cells per well for cell growth assay. In the migration assay, the confluent cell layers were pretreated with Mitomycin C (sigma aldrich, St. Louis, MO) for two hours, then scratched by using Essen wound maker. The plates were placed into IncuCyte (Sartorius) and imaged every two hours. The photographs were analyzed by using the IncuCyte software (Sartorius). Cell proliferation was also assessed by 5-ethynyl-2’-deoxyuridine (EdU) incorporation assay using Click-iT™ EdU Alexa Fluor™ 647 Flow Cytometry Assay Kit (Thermo Fisher Scientific) according to the manufacturer’s instructions.

### RNA extraction and qRT-PCR

Before RNA extraction, skin and wound biopsies were homogenized using TissueLyser LT (Qiagen, Hilden, Germany). Total RNA was extracted from human tissues and cells using the Trizol reagent (ThermoFisher Scientific). Reverse transcription was performed using the RevertAid First Strand cDNA Synthesis Kit (ThermoFisher Scientific). Gene expression was determined by TaqMan expression assays (ThermoFisher Scientific) or by SYBR Green expression assays (ThermoFisher Scientific) and normalized based on the values of the housekeeping gene *B2M* or *GAPDH*. Information for all the primers used in this study is listed in Supplementary Table 4.

### Magnetic cell separation of CD45-epidermal cells

After being washed in PBS, the skin and wound tissues were incubated in dispase II (5 U/mL, ThermoFisher Scientific) at 4°C overnight, and the epidermis and the dermis were separated. The epidermis was cut into small pieces and digested in 0.025% Trypsin/EDTA Solution at 37°C for 15 minutes. CD45- and CD45+ cells were separated using CD45 Microbeads with MACS MS magnetic columns according to the manufacturer’s instructions (Miltenyi Biotec, Bergisch Gladbach, Germany).

### Single-cell RNA sequencing

Skin and wounds were incubated in dispase II (5 U/mL, ThermoFisher Scientific) at 4°C overnight, and the epidermis and the dermis were separated. Single dermal cells were obtained using a whole skin dissociation kit (Miltenyi Biotec). Single epidermal cells were collected by incubating in 0.025% Trypsin-EDTA at 37°C for 15-20 minutes. After removing red blood cells, an equal number of epidermal and dermal cells were mixed. The dead cells were removed using the MACS Dead cell removal kit (Miltenyi Biotec). Single cells were loaded onto each channel of the Single Cell chip (10x Genomics, Pleasanton, CA) before droplet encapsulation on the Chromium Controller. Sequencing libraries were generated using the library construction kit (10x Genomics) following the manufacturer’s protocol. Libraries were sequenced on Illumina NovaSeq 6000 S4 v1.5. Raw sequencing data were processed with the cellranger pipeline (10X Genomics, version 5.0.1) and mapped to the hg38 reference genome to generate matrices of gene counts by cell barcodes.

Gene-barcode counts matrices were analyzed with the Seurat R package (version 4.0.6)^66^. Genes expressed in less than ten cells and cells with < 500 genes detected, < 1000 UMI counts, or > 20% mitochondrial gene mapped reads were filtered away from downstream analysis. The Seurat object of each sample was normalized and scaled with R package sctransform^67^. PCA was performed on the top 4000 variable genes using RunPCA, and the first 40 principal components (PCs) were used in the function of RunHarmony from the Harmony package^68^ to remove potential confounding factors among samples processed in different libraries. UMAP plots were generated using the RunUMAP function with the first 40 harmonies. The clusters were determined by using the FindNeighbors and FindClusters functions with a resolution of 0.8.

### Bulk RNA-sequencing

Ribosomal RNAs (rRNA) were removed using the Epicentre Ribo-zero® rRNA Removal Kit (Epicentre, Madison, WI). For the whole biopsies, strand-specific total-transcriptome RNA sequencing libraries were prepared by incorporating dUTPs in the second-strand synthesis using NEB Next® Ultra^TM^ Directional RNA Library Prep Kit for Illumina® (NEB, Ipswich, MA). The isolated cell RNA-seq libraries were prepared using the Ovation® SoLo® RNA-Seq library preparation kit (Tecan, Männedorf, Switzerland). The libraries were sequenced on the Illumina Hiseq 4000 platform (Illumina, San Diego, CA).

Raw reads of RNA sequencing were first cleaned by using Trimmomatic v0.36 software^69^. Cleaned reads were mapped to the human reference genome (GRCh38.p12) and the comprehensive gene annotation file (GENCODEv31) using STAR v2.7.1a^70^. Gene expression was then quantified by calculating unique mapped fragments to exons using the Subread package^71^. The differential expression analysis was performed using DESeq2^72^. The differentially expressed RNAs were defined as P < 0.05, and |Fold change| >= 2. Raw counts for each gene were normalized to fragments per kilobase of a transcript, per million mapped reads (FPKM)-like values.

### *In situ* hybridization

*HOXC13-AS* probe (Cat. No. 486901) was designed and synthesized by Advanced Cell Diagnostics (ACD, Silicon Valley, CA). Tissues and cells were prepared by following the manufacturer’s instructions. After fixation and dehydration with 50%, 70%, and 100% ethanol, the slides were incubated with Protease IV (ACD) at room temperature for 30 minutes. Then the slides were incubated with *HOXC13-AS* probes for two hours at 40 °C in HybEZ™ II Hybridization System using RNAscope® Multiplex Fluorescent Reagent Kit v2 (ACD). The hybridization signals were amplified via sequential hybridization of amplifiers and label probes. *HOXC13-AS* signals were visualized on Zeiss LSM800 confocal microscope (Oberkochen, Germany) and analyzed with ImageJ software (National Institutes of Health).

### Western blotting

Keratinocyte cell lysates were extracted using radioimmunoprecipitation assay (RIPA) buffer complemented with protease inhibitor and phosphatase inhibitor. The total protein was separated in TGX™ precast protein gels (Bio-Rad, Hercules, CA) and transferred to a nitrocellulose membrane. Blots were probed with KRT10 antibody (ab76318, 1:1000 dilution, Abcam, Cambridge, United Kingdom) or antibodies for ER stress markers (9956, dilution following manufacturer’s instructions, CST, Danvers, MA). Next, the blots were incubated with anti-mouse (P0447, 1:1000 dilution, DAKO, Denmark) or anti-rabbit (P0448, 1:2000 dilution, DAKO) HRP conjugated secondary antibodies. β-actin levels were used as an internal control in each sample (A3854, 1:20,000 dilution, Sigma-Aldrich, St. Louis, MO). Protein bands were quantified with ImageJ and Image lab (Bio-Rad).

### Suspension-induced keratinocyte differentiation

Keratinocytes differentiate in suspension, as described previously^34^. We suspended keratinocytes at a concentration of 10^5^ cells/mL in the EpiLife medium containing 1.45% methylcellulose (Sigma-Aldrich) in 24-well plates at 37°C. At different time points after cell suspension, the methylcellulose was diluted with PBS and the cells were recovered by centrifugation.

### Immunofluorescence staining

Cultured cells were fixed with 4% paraformaldehyde for 15 minutes at room temperature and permeabilized with 0.1% Triton X-100. The paraffin-embedded organotypic epidermis was deparaffinized and rehydrated before staining. Cells and tissues were blocked by 5% bovine serum albumin (BSA) in PBS for 30 minutes at room temperature, followed by immunostaining with anti-Ki67 (9129, 1:400 dilution, CST), anti-GM130 (PA1-077, 1:500 dilution, ThermoFisher Scientific), anti-KRT10 (M7002, 1:200 dilution, DAKO), or anti-FLG antibodies (MA513440, 1:100 dilution, ThermoFisher Scientific) at 4℃ overnight. On the next day, the sections were washed three times with PBS and then incubated with donkey anti-mouse or anti-rabbit secondary antibody conjugated with Alexa Fluor™ 488 (A-21202 and A-21206, 1:1000 dilution, Invitrogen) in the dark at room temperature for 40 minutes. After being washed with PBS three times, the sections were mounted with the ProLong™ Diamond Antifade Mountant with DAPI (ThermoFisher Scientific). Samples were visualized by Zeiss LSM800 confocal microscopy and analyzed with ImageJ software.

### Gene expression microarray and analysis

Keratinocytes were transfected with either *HOXC13-AS* siRNA or control siRNA for 24 hours (n=3 per group) and transcriptomic profiling was performed using human Clariom™ S assays (ThermoFisher Scientific) at the core facility for Bioinformatics and Expression Analysis (BEA) at Karolinska Institute. In brief, total RNA was extracted using the miRNeasy Mini Kit (Qiagen), and RNA quality and quantity were determined using Agilent 2200 Tapestation with RNA ScreenTape (Agilent, Santa Clara, CA) and Nanodrop 1000 (ThermoFisher Scientific). 150 ng of total RNA was used to prepare cDNA following the GeneChip WT PLUS Reagent Kit labeling protocol (ThermoFisher Scientific). Standardized array processing procedures recommended by Affymetrix, including hybridization, fluidics processing, and scanning, were used. Genes showing at least 2-fold change and with a P value less than 0.05 were considered differentially expressed. Gene set enrichment analysis (GSEA) was performed using a public software from Broad Institute^33^. MetaCore software (Thomson Reuters, Toronto, Canada) was used for Gene Ontology (GO) analysis. Heatmaps were generated with the Multiple Experiment Viewer software.

### Organotypic epidermis culture

The organotypic epidermis was cultured as described previously^16^. 1.55 × 10^5^ human primary keratinocytes were resuspended in a CnT-PR medium (CELLnTEC, Bern, Switzerland) and placed onto 0.47 cm^2^ inserts with 0.4 μm pore size (Nunc, Roskilde, Denmark). Six inserts were put into a 60 mm cell culture dish filled with the CnT-PR medium. When cells reached confluency, the medium inside and outside the insert was replaced with a CnT-PR-3D medium (CELLnTEC). Twenty-four hours later, the medium inside was removed, and the surface of the inserts remained dry for the rest of the cultivation. Medium outside was changed every two days. Inserts were collected on day 12 after airlifting and followed by fixation and embedding.

### Cellular fractionation

Nuclear and cytoplasmic fractions were separated by PARIS kit following manufacturer’s instructions (ThermoFisher Scientific). RNA was extracted from these fractions by using Trizol (ThermoFisher Scientific) and qRT-PCR was performed to analyze the *MALAT1*, *B2M*, *GAPDH,* and *HOXC13-AS* expression.

### *HOXC13-AS* overexpression construct

For the generation of the *HOXC13-AS* expression vector, the *HOXC13-AS* fragment in pEX-A258 was obtained from Eurofins Genomics (Ebersberg, Germany) and cloned into pcDNA3.1(+) vector (Invitrogen, Carlsbad, CA). The construct was further sequenced to verify the orientation and integrity of the ligated *HOXC13-AS* insert.

### RNA pulldown

Full-length *HOXC13-AS* was transcribed *in vitro* using the T7 MEGAscript kit (ThermoFisher Scientific) and labeled using the Pierce RNA 3′ End Desthiobiotinylation Kit (ThermoFisher Scientific). *HOXC13-AS* pulldown assay was performed using the Pierce Magnetic RNA-Protein Pull-Down Kit (ThermoFisher Scientific) according to the manufacturer’s instructions. Briefly, 50 pmol of desthiobiotinylated *HOXC13-AS* was mixed with magnetic beads and incubated with 200 µg of protein lysate from human adult epidermal keratinocytes (ThermoFisher Scientific). Poly(A)_25_ RNA, which was included in the RNA-Protein Pull-Down kit, was used as a negative control. The beads were washed, and RNA-bound proteins were eluted by boiling the beads in SDS buffer. Eluted protein samples were run on the Mini-PROTEAN TGX gels (Bio-Rad) followed by silver staining. Specific bands were excised and analyzed by mass spectrometry at the Proteomics Biomedicum core facility, Karolinska Institute.

### In-gel protein digestion and mass spectrometry

Protein bands were excised manually from gels and in-gel digested using a MassPREP robotic protein-handling system (Waters, Millford, MA). Gel pieces were distained following the manufacturer’s description. Proteins then were reduced with 10 mM DTT in 100 mM Ambic for 30 min at 40°C and alkylated with 55 mM iodoacetamide in 100 mM Ambic for 20 min at 40°C followed by digestion with 0.3 μg trypsin (sequence grade, Promega, Madison, WI) in 50 mM Ambic for 5 hours at 40°C. The tryptic peptides were extracted with 1% formic acid in 2% acetonitrile, followed by 50% acetonitrile twice. The liquid was evaporated to dryness on a vacuum concentrator (Eppendorf).

The reconstituted peptides in solvent A (2% acetonitrile, 0.1% formic acid) were separated on a 50 cm long EASY-spray column (ThermoFisher Scientific) connected to an Ultimate-3000 nano-LC system (ThermoFisher Scientific) using a 60 min gradient from 4-26% of solvent B (98% acetonitrile, 0.1% formic acid) in 55 min and up to 95% of solvent B in 5 min at a flow rate of 300 nL/min. Mass spectra were acquired on a Q Exactive HF hybrid Orbitrap mass spectrometer (ThermoFisher Scientific) in m/z 375 to 1500 at resolution of R=120,000 (at m/z 200) for full mass, followed by data-dependent HCD fragmentations from most intense precursor ions with a charge state 2+ to 7+. The tandem mass spectra were acquired with a resolution of R=30,000, targeting 2×10^5^ ions, setting isolation width to m/z 1.4 and normalized collision energy to 28%.

Acquired raw data files were analyzed using the Mascot Server v.2.5.1 (Matrix Science Ltd., UK) and searched against SwissProt protein databases (20,368 human entries). Maximum of two missed cleavage sites were allowed for trypsin, while setting the precursor and the fragment ion mass tolerance to 10 ppm and 0.02 Da, respectively. Dynamic modifications of oxidation on methionine, deamidation of asparagine and glutamine and acetylation of N-termini were set. Initial search results were filtered with 5% FDR using Percolator to recalculate Mascot scores. Protein identifications were accepted if they could be established at greater than 96.0% probability and contained at least 2 identified peptides. Proteins that contained similar peptides and could not be differentiated based on MS/MS analysis alone were grouped to satisfy the principles of parsimony.

### RNA immunoprecipitation

RNA immunoprecipitation (RIP) was performed using the Magna RIP RNA-Binding Protein Immunoprecipitation Kit (Millipore, Burlington, MA). Briefly, human primary keratinocytes were lysed in RIP lysis buffer, and then 100 μl of cell lysate was incubated with an anti-COPA antibody (sc-398099, Santa Cruz Biotechnology, Dallas, TX). Mouse IgG, instead of anti-COPA antibody, was used as a negative control. The immunoprecipitated RNA was extracted and analyzed using RT-PCR followed by electrophoresis.

### Statistics

Data analysis was performed by using GraphPad Prism Version 7 (Dotmatics, San Diego, CA). Statistical significance among the two groups was determined by a two-tailed Student’s t-test or Mann-Whitney Test. The significance among multiple groups was determined by one-way ANOVA with Bonferroni posttest. Pearson’s correlation test was performed on log2-transformed data. *P* < 0.05 was considered statistically significant.

### Schematics

Schematic cartoons in 1a, 5a, 6b, 6i, 7i, and Supplementary Fig. 3k were created with BioRender.com.

### Data availability

Raw data of long RNA sequencing and microarray performed for this study have been deposited to NCBI’s Gene Expression Omnibus (GEO) database under the accession number GSE174661 and GSE206103, respectively.

## ACKNOWLEDGEMENT

We express our gratitude to all the tissue donors participating in this study. We thank the Microarray core facility at Novum, BEA, which is supported by the board of research at Karolinska Institute and the research committee at the Karolinska hospital. In-gel digestion, peptide extraction, mass spectrometric analysis, and database searches for protein identification were carried out at the Proteomics Biomedicum, Karolinska Institute, Stockholm. The computations/data handling was enabled by resources in the projects sens2020010 and 2021/22-701 provided by the Swedish National Infrastructure for Computing (SNIC) at UPPMAX, partially funded by the Swedish Research Council through grant agreement no. 2018-05973. This work was supported by Swedish Research Council (Vetenskapsradet, 2020-01400), Ragnar Söderbergs Foundation (M31/15), Welander and Finsens Foundation (Hudfonden), LEO foundation, Cancerfonden, Ming Wai Lau Centre for Reparative Medicine, and Karolinska Institute.

## Author contributions

NXL and LZ conceived and designed the study. PS collected most clinical samples with the assistance of MAT and XB. LZ, MP, DL, and XB performed the experiments. ZL and LZ carried out bioinformatics analysis. LZ, MP, and NXL contributed to data analyses and interpretation. LZ, MP, and NXL wrote the manuscript, which was commented on by all the authors.

## Competing interests

The authors declare no competing interests.

## Additional information

The supplementary materials are available.

Correspondence and requests for materials should be addressed to Ning Xu Landén.

## Notes

### Competing Interest Statement

The authors have declared no competing interest.

