## Supplementary files for "Human skin specific long noncoding RNA *HOXC13-AS* regulates epidermal differentiation by interfering with Golgi-ER retrograde transport"

### This PDF file includes:

Supplementary Figures 1–5

Supplementary Tables 1–4

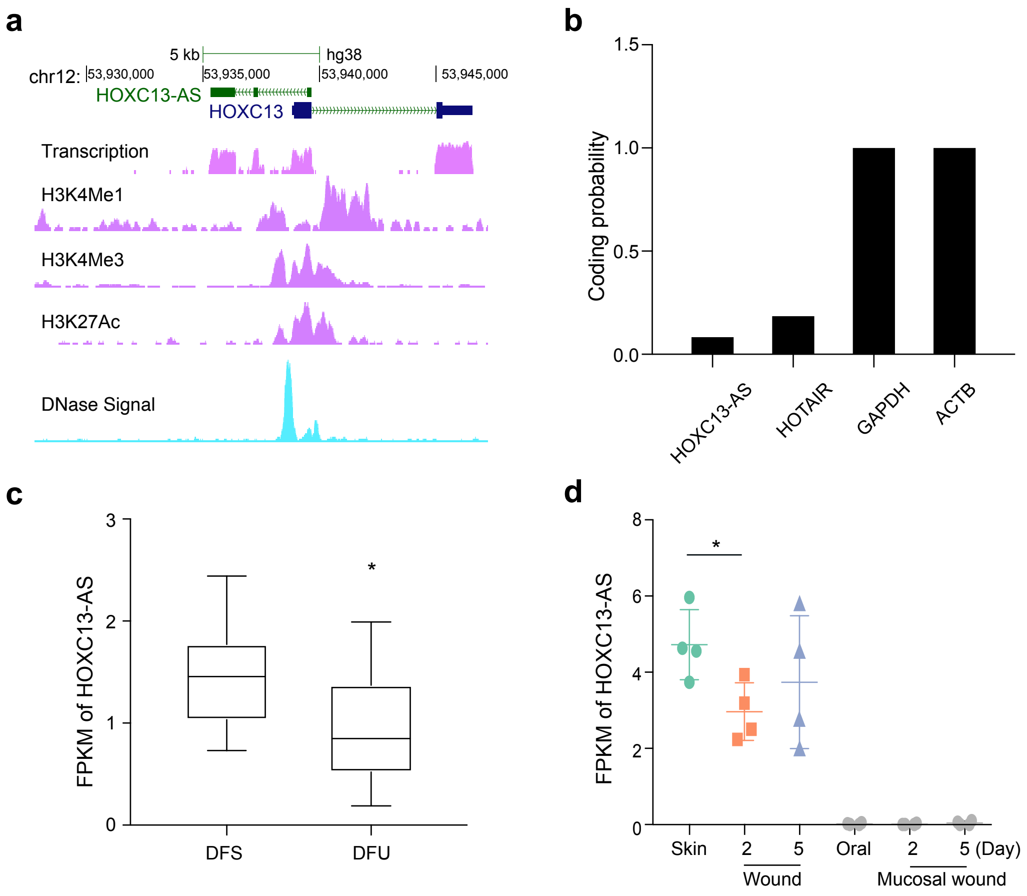


**Supplementary Figure 1 Characterization of *HOXC13-AS.* a** UCSC genome browser tracks depicting transcription, DNase hypersensitivity regions, and histone modifications (H3K4me1, H3K4me3, H3K27ac) in *HOXC13-AS* and *HOXC13* loci. **b** Bar graph representing coding potential scores obtained from Coding Potential Calculator 2 for coding (*GAPDH* and *ACTB*) and non-coding (*HOXC13-AS* and *HOTAIR*) genes. Querying *HOXC13-AS* expression in the published RNA-seq datasets of human wounds: diabetic foot ulcers (DFU) vs. diabetic foot skin (DFS) (GSE134431) **c**; day -2 and day-5 skin wounds vs. day -2 and day-5 mucosal wounds (GSE97615) **d**. Data are normalized as Fragments per kilobase of a transcript, per million mapped reads (FPKM).

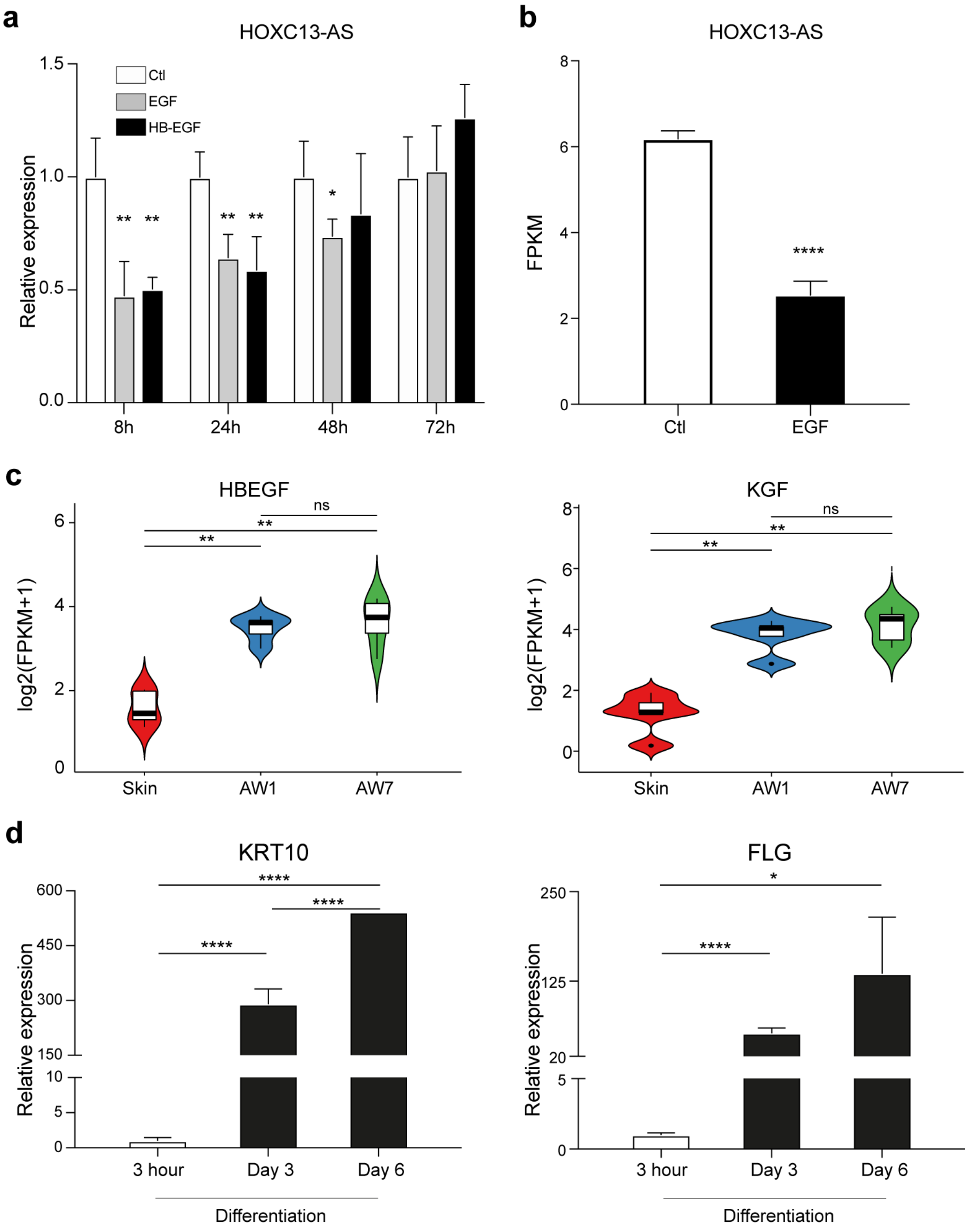


**Supplementary Figure 2 *HOXC13-AS* expression in keratinocytes is oppositely regulated by growth and differentiation signals. a** qRT-PCR analysis of *HOXC13-AS* expression in human primary keratinocytes treated with EGF or HBEGF for 8-72 hours (n = 4). **b** Querying *HOXC13-AS* expression in the published RNA-seq datasets of epidermal stem cells treated with EGF (GSE156089). **c** RNA-seq analysis of *HBEGF* and *KGF* expression in human skin and day-1 and day-7 acute wounds. **d** qRT-PCR analysis of *KRT10* and *FLG* expression in calcium-induced differentiated keratinocytes (n = 4). **P* < 0.05; ***P* < 0.01 and *****P* < 0.0001 by unpaired two-tailed Student's t test **a**, **b**, **d**, or Mann Whitney test **c**. Data are presented as mean ± SD or violin plots.

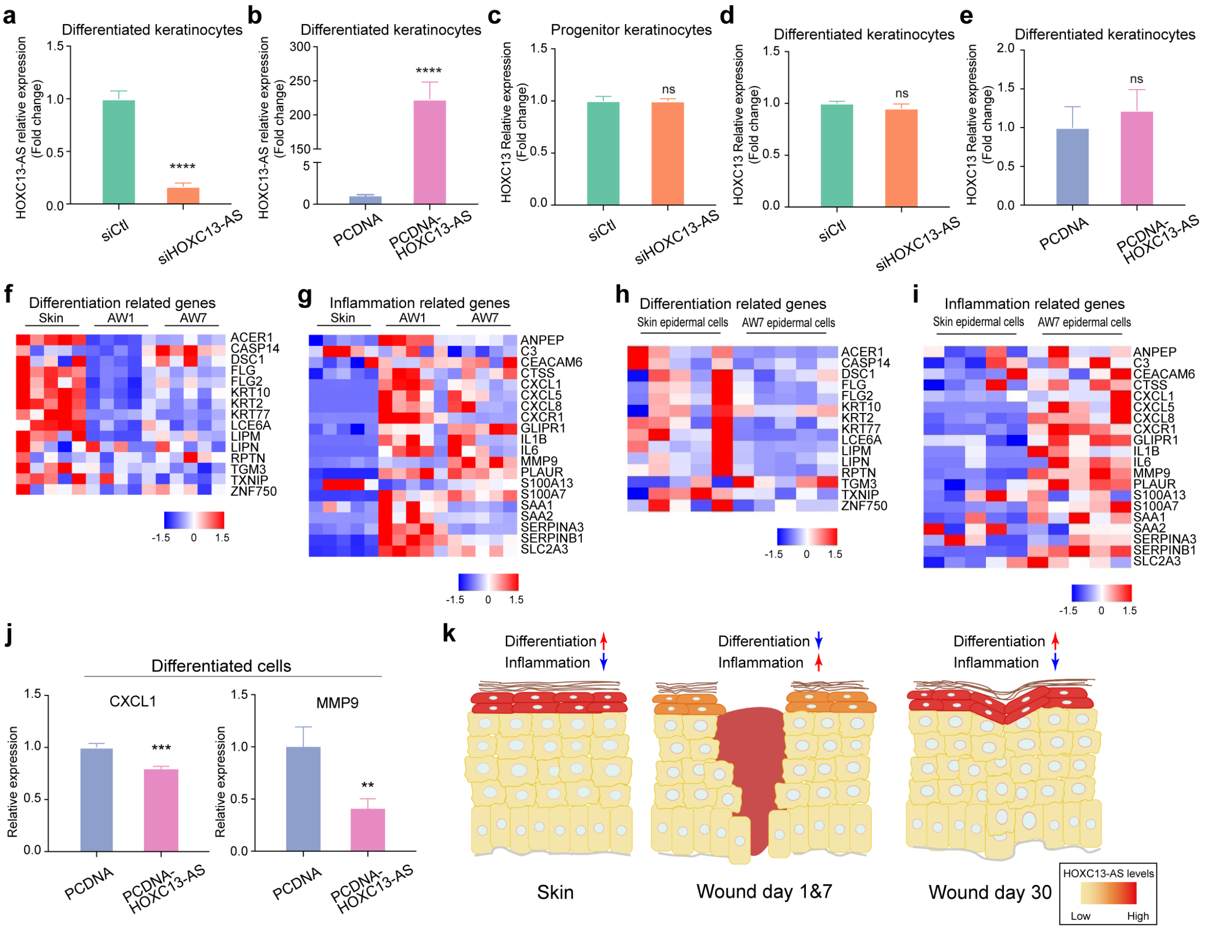


**Supplementary Figure 3 *HOXC13-AS* regulates keratinocyte differentiation and inflammatory response.** qRT-PCR analysis of *HOXC13-AS* expression in keratinocyte transfected with *HOXC13-AS* siRNA pool **a**, or pcDNA-*HOXC13-AS* **b** (n = 4). qRT-PCR analysis of *HOXC13* expression in keratinocyte transfected with HOXC13-AS siRNA pool **c**, **d**, or pcDNA-HOXC13-AS **e** in progenitor or differentiated keratinocytes. Heatmaps showing the expression of HOXC13-AS regulated genes in the skin and wound tissue biopsies **f**, **g** and the isolated epidermal cells **h**, **i** analyzed by RNA-seq. **j** qRT-PCR analysis of CXCL1 and MMP9 in differentiated keratinocytes with *HOXC13-AS* overexpression (n = 4). **k** Schematic representation of the changes of *HOXC13-AS* expression and its regulated biological processes during human skin wound healing. ns: not significant, **$P < 0.01$, ***$P < 0.001$ and ****$P < 0.0001$ by unpaired two-tailed Student's t test **a-e**, **j**. Data are presented as mean ± SD.

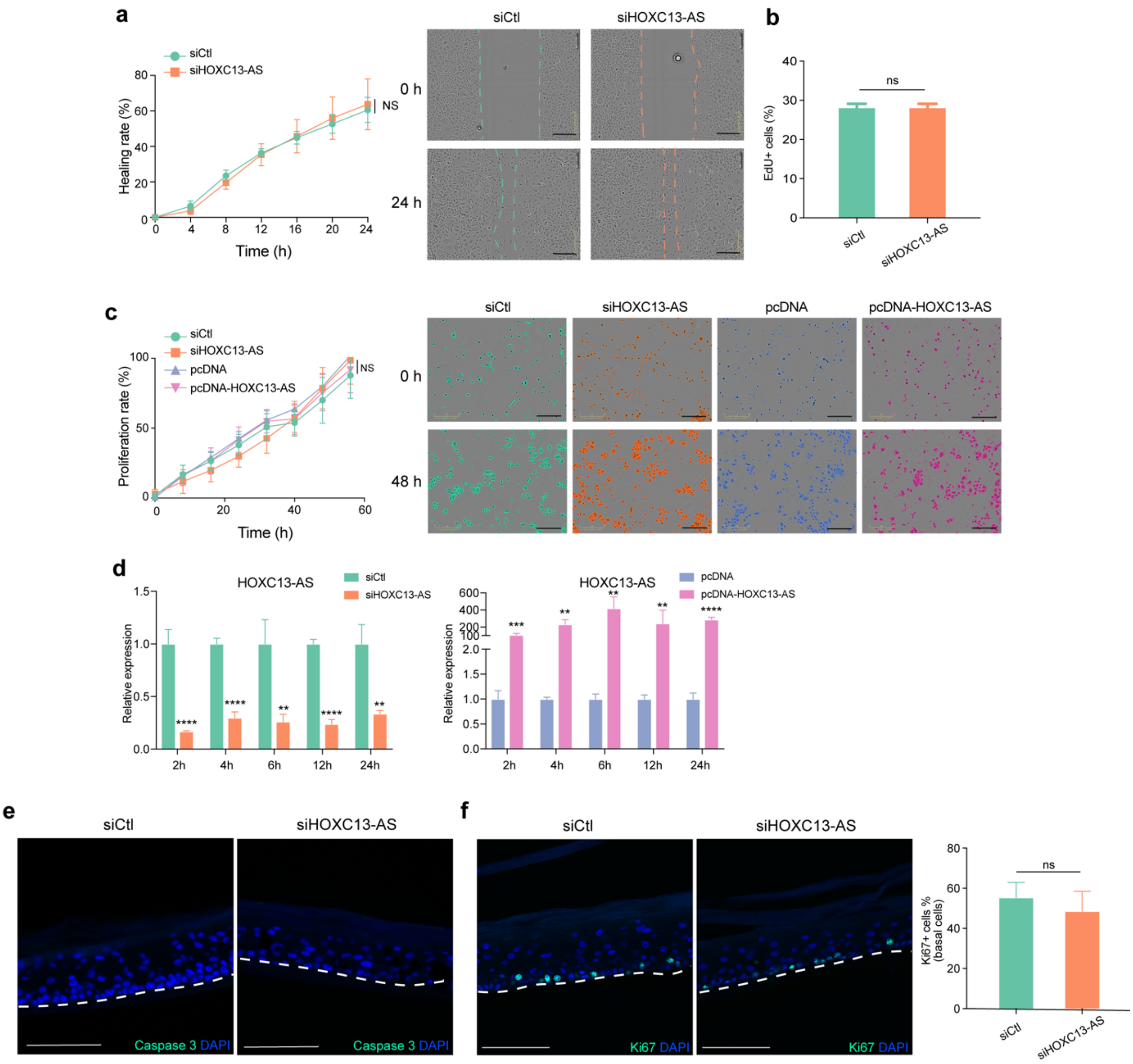


**Supplementary Figure 4** ***HOXC13-AS*** **does not affect keratinocyte growth, migration, or apoptosis. a** The migration of keratinocytes with *HOXC13-AS* knockdown (n = 8) was analyzed using a live cell imaging system. Representative photographs of cells at 0 and 24 hours are shown (Scale bar, 300 μm). **b** 5-ethynyl-2'-deoxyuridine (EdU) incorporation assay of keratinocytes with *HOXC13-AS* silencing (n = 4). **c** The growth of keratinocytes with *HOXC13-AS* knockdown or overexpression (n = 3) was analyzed using a live cell imaging system. Representative photographs of cells at 0 and 96 hours are shown (Scale bar, 300 μm.). **d** qRT-PCR analysis of *HOXC13-AS* expression in keratinocytes with *HOXC13-AS* knockdown or overexpression followed by suspension induction for 2-24 hours (n = 3). Immunofluorescent staining of caspase 3 **e** and Ki67 **f** in organotypic human epidermal tissues with *HOXC13-AS* knockdown (scale bar = 100 μm, n = 3). ns: not significant, $**P < 0.01$, $***P < 0.001$, and $****P < 0.0001$ by unpaired two-tailed Student's t test **b**, **d**, **f**, or one-way ANOVA **a**, **c**. Data are presented as mean ± SD.

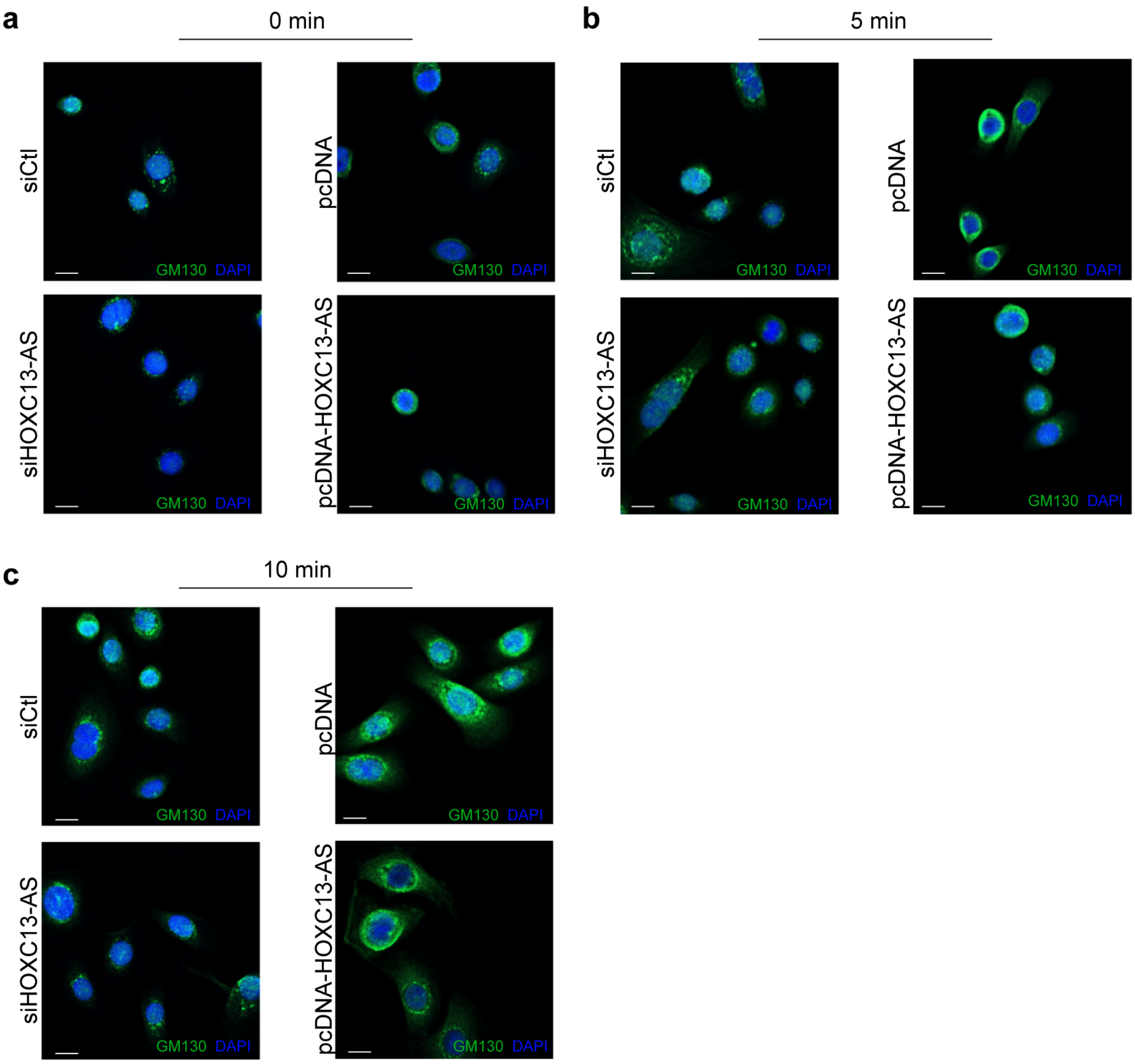


**Supplementary Figure 5** Representative photograph of GM130 immunofluorescent staining in keratinocytes transfected with *HOXC13-AS* siRNA pool, pcDNA-*HOXC13-AS,* or respective controls, and treated with Brefeldin A for 0 minute **a**, 5 minutes **b,** and 10 minutes **c**. Cell nuclei were co-stained with DAPI. Scale bar = 10 μm

**Supplementary Table 1. Donor information**

| Donor | Sex | Age (years) | Experiments |
|---|---|---|---|
| 1 | F | 66 | RNA sequencing |
| 2 | M | 69 | RNA sequencing |
| 3 | F | 67 | RNA sequencing |
| 4 | M | 69 | RNA sequencing, qRT-PCR |
| 5 | F | 64 | RNA sequencing, qRT-PCR |
| 6 | F | 60 | qRT-PCR |
| 7 | F | 66 | qRT-PCR |
| 8 | F | 60 | qRT-PCR |
| 9 | F | 67 | qRT-PCR |
| 10 | F | 65 | qRT-PCR |
| 11 | M | 26 | Cell isolation, RNA sequencing |
| 12 | F | 30 | Cell isolation, RNA sequencing |
| 13 | M | 45 | Cell isolation, RNA sequencing |
| 14 | F | 43 | Cell isolation, RNA sequencing |
| 15 | M | 22 | Cell isolation, RNA sequencing |
| 16 | F | 48 | Cell isolation, qRT-PCR |
| 17 | M | 27 | Cell isolation, qRT-PCR |
| 18 | F | 24 | Cell isolation, qRT-PCR |
| 19 | M | 24 | Cell isolation, qRT-PCR |
| 20 | F | 46 | Cell isolation, qRT-PCR |
| 21 | F | 24 | ISH |
| 22 | F | 22 | ISH, scRNAseq |
| 23 | M | 29 | scRNAseq |
| 24 | M | 24 | scRNAseq |

M, male; F, female

**Supplementary Table 2. List of cytokines/growth factors used in the study**

| Cytokines/Growth factors | Cat.no | Concentration | Vendor |
|---|---|---|---|
| EGF | 11343406 | 20ng/mL | ImmunoTools |
| FGF-2 | 11343625 | 30ng/mL | ImmunoTools |
| GM-CSF | 11343123 | 50ng/mL | ImmunoTools |
| HB-EGF | 259-HE-050 | 20ng/mL | ImmunoTools |
| IGF-1 | 11343314 | 20ng/mL | ImmunoTools |
| IL-36a | 11340362 | 10ng/mL | ImmunoTools |
| IL-6 | 11340060 | 50ng/mL | ImmunoTools |
| IL1α | 11349013 | 20ng/mL | ImmunoTools |
| IL23 | 11340232 | 10ng/mL | ImmunoTools |
| KGF | 11343653 | 20ng/mL | R&D Systems |
| MCP-1 | 11343380 | 10ng/mL | ImmunoTools |
| TGFB1 | 11343161 | 10ng/mL | ImmunoTools |
| TGFB3 | 11344482 | 10ng/mL | ImmunoTools |
| TNF-a | 210-TA | 50ng/mL | R&D Systems |
| VEGF | 11343663 | 20ng/mL | ImmunoTools |

**Supplementary Table 3**. **List of siRNA pools used in the study**

| siRNAs | Cat.no/Vendor |
|---|---|
| siHOXC13-AS pool | R-188045-00-0005, Dharmacon$^{TM}$, Lafayette, CO |
| Lincode Non-targeting pool | D-001320-10-05, Dharmacon$^{TM}$ |
| Non-Targeting pool | D-001810-10-05, Dharmacon$^{TM}$ |
| EIF3A | L-0195300-0005, Dharmacon$^{TM}$ |
| HDLBP | L-019956-00-0005, Dharmacon$^{TM}$ |
| COPA | L-011835-00-0005, Dharmacon$^{TM}$ |

**Supplementary Table 4**. **List of primers used in the study**

| Gene | Sequence/Cat.no/Vendor |
|---|---|
| HOXC13-AS | F 5' CGGAGAAAGACGGAGCATTTA<br>R 5' CAGAGTTGAACTTAGCCGAGAG, IDT |
| KRT10 | Hs.PT.58.38635764, IDT |
| FLG | F 5' ATGAGCAGGCACGAAACA<br>R 5' CCTGAGTGTCCAGACCTATCTA, IDT |
| IVL | Hs.PT.58.39460547, IDT |
| MALAT1 | F 5' GATCTAGCACAGACCCTTCAC<br>R 5'CGACACCATCGTTACCTTGA, IDT |
| sXBP1 | 356738665 Rxn Ready Primer Pool, IDT |
| tXBP1 | 356738667 Rxn Ready Primer Pool, IDT |
| ATF4 | 356738668 Rxn Ready Primer Pool, IDT |
| CHOP | 356738669 Rxn Ready Primer Pool, IDT |
| BiP | 356738670 Rxn Ready Primer Pool, IDT |
| EDEM | 356738672 Rxn Ready Primer Pool, IDT |
| IL6 | Hs.PT.5840226675, IDT |
| IL8 | Hs.PT.5839926886.g, IDT |
| IL1B | Hs.PT.581518186, IDT |
| B2M | F 5' AAGTGGGATCGAGACATGTAAG<br>R 5' GGAGACAGCACTCAAAGTAGAA, IDT |
| GAPDH | HS.PT.39A.22214836(1), IDT<br>F 5' GGTGTGAACCATGAGAAGTATGA<br>R 5' GAGTCCTTCCACGATACCAAAG, IDT |
